# Large phenological advances and delays over 124 years of climate change alter co-flowering among North American *Viola*

**DOI:** 10.64898/2026.06.27.734973

**Authors:** Caroline Edwards, Leonie C. Moyle

## Abstract

Shifts in flowering phenology are one of the most well studied plant responses to global climate change. Many studies have documented these shifts and their drivers, including some that describe altered patterns of co-flowering among taxonomically diverse species within communities. In comparison, few analyses have examined systematic changes in co-flowering between closely related, interfertile species, where co-flowering can have unique evolutionary consequences. To address such shifts in co-flowering among close relatives, we investigate phenological responses to climate change and its effect on patterns of co-flowering over the past 124 years in 52 species of North America violets (*Viola*). This genus has many co-occurring species that reproductively interact via shared pollinators and hybridization. We use ∼14,000 herbarium records along with environmental and species trait data to model the magnitude of recent flowering phenology shifts, environmental variables and/or species traits associated with these shifts, and resulting changes in co-flowering among species. While both the magnitude and direction of phenological shifts varied among *Viola* species, nearly half (25/52) show significant changes in flowering day. Regardless of whether flowering was advanced or delayed, flowering date was most consistently associated with local mean temperature. Of six species-level traits, geographical region also significantly predicted flowering shifts, consistent with environment and geography together explaining broad phenological responses across this group. These shifts have produced significant changes to pairwise patterns of co-flowering among species—ranging from a 59 day increase in co-flowering to complete loss of co-flowering overlap. Sympatric pairs specifically have experienced both increases and decreases in co-flowering, within a significant geographic pattern of increased co-flowering mainly in eastern US and decreased co-flowering common in western US. Because *Viola* species are generalist pollinated and already known to hybridize, these new co-flowering patterns could further undermine reproductive barriers among species in this genus.

## Introduction

Climate change is altering species’ environments, with complex consequences for both individual species and for species interactions. For plants, one common phenotypic response to climate change is a shift in the timing of key life events, including the timing of flowering (Parmesan & Yohe, 2006; Menzel et al., 2006; Williamson et al., 2025). Climate-mediated flowering shifts are important for individual species’ responses to changing environmental conditions, but can also impact reproductive interactions among species by altering which species co-flower and the duration of their co-flowering period (CaraDonna et al., 2014; Austin et al., 2024; Forrest et al., 2010). This altered co-flowering could affect pollinator visitation rates via competition or facilitation (Faust & Iler, 2022), as well as the likelihood of reproductive interference or hybridization among heterospecifics—changes that could be particularly important for closely-related, interfertile species for whom flowering phenology can be an important reproductive isolating mechanism (Franks & Weis, 2009). Understanding how and why patterns of flowering and co-flowering have changed over recent time, especially among close relatives, is therefore critical to understanding the potential evolutionary consequences of flowering phenology shifts due to climate change.

For individual species, shifts in flowering phenology have been associated with changes in environmental variables such as temperature (Menzel et al., 2006; Augspurger et al., 2020; Williams et al., 2021), precipitation (König et al., 2018) and the interaction of these factors (Matthews & Mazer, 2016). Increased temperature is widely associated with advanced flowering, especially for spring flowering species in temperate regions (Miller-Rushing & Primack, 2008; Park et al., 2018). Precipitation has a more variable effect on flowering shifts, but decreased precipitation has been associated with advanced flowering in arid (Crimmins et al., 2010) and temperate regions (König et al., 2018). Because environmental conditions are spatially structured, species responses also can vary among geographic regions (Templ et al., 2017; Kopp et al., 2020; Williamson et al., 2025). Moreover, even within a geographic region or across similar environments, responses to changing environmental conditions can be species-specific (Park et al., 2018).

One important factor that might shape these species-specific phenological responses is variation due to species’ traits (Pareja-Bonilla et al., 2026), including origin status (native or introduced) and life history (annual or perennial). For instance, in response to increasing temperatures, introduced species have been shown to advance their flowering more than native species (Calinger et al., 2013). Annual herbs have advanced flowering more than perennials across multiple genera (Fitter & Fitter, 2002; Miller-Rushing & Primack, 2008) and geographic locations (König et al., 2018), potentially because annuals have a shorter generation time and therefore faster evolutionary rates (König et al., 2018). Numerous other species properties have been proposed to explain variable responses to climate change, including differences in flowering season (Calinger et al., 2013; Fitter & Fitter 2002), reliance on external pollinators via pollination mode and mating system (Molnar et al., 2012; König et al., 2018), and genetic variation (Doi et al., 2010).

An important consequence of heterogeneous, species-specific phenological responses to changing climate is the emergence of new patterns of co-flowering among co-occurring species. Such changes in co-flowering (“phenological reassembly”; Theobald et al., 2017) have been evaluated among diverse plant species within single communities (e.g. CaraDonna et al., 2014; Austin et al., 2024; Pareja-Bonilla et al., 2025), and across communities at larger spatial scales (e.g. Fisogni et al., 2022; Ramirez-Parada et al., 2025). In each case, recent climate change has been accompanied by altered co-flowering patterns, but the relative fraction of increased vs decreased co-flowering species (and the magnitude of change in co-flowering) varies among communities. Moreover, while these changes in co-flowering could potentially have important evolutionary consequences, including altered rates of pollinator visitation and heterospecific pollen transfer (Sargent & Ackerly, 2009), few studies focus on closely-related congeners that are uniquely vulnerable to additional effects such as species hybridization. In one such exception, Carsadden et al. (2022) investigated climate-associated phenological shifts in two sympatric and hybridizing *Potentilla* species, but found no change in flowering overlap due to similar phenological responses in both species. In general, however, the consequences of co-flowering shifts for changing reproductive contact between congeneric community members remain poorly understood.

One approach to assessing evidence for or against such consequences is to examine phenological shifts in many closely related species across a contiguous geographic region. In addition to quantifying individual species responses to altered climate, this approach provides the power to examine general relationships between the magnitude and direction of observed phenological shifts and species’ variation in environmental and biological factors. In combination with information about species’ ranges and their pairwise range overlap, these data can also be used to assess whether phenological shifts have altered temporal patterns of species co-flowering. Here we take this multispecies approach to investigate the degree to which *Viola* species in North America have shifted their flowering phenology over the past 124 years, and to explore possible causes and consequences of this trait change. We draw on geolocation, environmental, and flowering time data from ∼14,000 herbarium records from 52 *Viola* species, as well as species trait data from the Flora of North America, to address these questions. First, we determine how flowering phenology has changed over the past 124 years for each species by quantifying the rate at which species are advancing or delaying flowering (days/year), using all flowering records. We also confirm that key environmental variables have changed over this period, at the geolocations where these 52 species have been recorded. Second, we analyze whether changes in two major environmental factors are associated with flowering phenology and whether these are consistent predictors of advanced or delayed flowering. Third, we test for associations between species’ rate of phenological shift since 1900 (in days/year) and species’ traits (such as range size and mating system) to understand potential drivers, or factors limiting, species responses to the changing environment. Finally, to connect these individual species responses with potential phenological reassembly among species, we examine how shifts in flowering phenology have changed patterns of co-flowering among species pairs, including those specifically in geographical sympatry.

## Materials and Methods

### System and study region

*Viola* is a speciose and geographically widespread genus of ∼600 species distributed worldwide, a majority of which are found in northern temperate zones. Violets are short-lived (annual or perennial), mostly herbaceous plants whose species occupy many habitat types including woodland understories, bogs, prairies and backyards (Little & McKinney, 1993). Their distributions vary from small, restricted ranges (e.g., *V. clauseniana*) to locally abundant and geographically widespread (e.g., *V. sororia*). In this study, we focus our analyses on the *Viola* species living in North America, for which there are high quality standardized climate and trait data available. Three non-native species from Europe grow commonly in North America, so we also included introduced North American populations of these species in our study.

*Viola* species have several distinctive characteristics that might influence their flowering phenological responses to recent climate change, and the ecological and evolutionary consequences of these responses. One such characteristic is a form of cleistogamy (self-pollination through non-opening flowers) called seasonal cleistogamy, in which an individual plant produces primarily outcrossing chasmogamous (CH) flowers in the spring, and then selfing cleistogamous (CL) flowers in late spring and summer. While seasonal cleistogamy (CHCL) is the most common reproductive strategy in *Viola*, several species in the genus produce only CH flowers (Little & McKinney, 1993). Whether CHCL mating system variation generates different phenological responses to recent climate change is unknown, but we hypothesize that CHCL species will shift flowering less than CH species, as species that are less reliant on external pollination have been shown to be less sensitive to changing climate (König et al., 2018). *Viola* species also vary in flower color—ranging from purple, to yellow, to white—and attract a wide range of insect pollinators including species in Hymonoptera (e.g. bumblebees, solitary bees), Syrphidae (hoverflies), Bombyliids (bee flies), and Lepidoptera (moths, butterflies) (Beattie, 1974). Most species are pollinated by several groups of insects and share pollinator groups. All species begin flowering in spring, but vary in duration of flowering window, with some species flowering until the end of spring while others continue flowering through the summer into early fall (Little & McKinney, 1993).

In addition, *Viola* species frequently co-occur with one or more congeners. Many species are also known to hybridize, suggesting that reproductive isolating barriers are often weak and potentially vulnerable to disruption in this group (Marcussen et al., 2022; Ballard et al., 2023). Records of hybrids extend back into the last century (Brainerd, 1904), indicating that this history of hybridization is not solely a product of recent climate change. However, whether the frequency of hybridization might change in the face of climate-mediated shifts in flowering phenology remains unknown.

### Flowering occurrence data

We used herbarium specimens to assess evidence for phenological shifts. We focused on digitized data from specimens that were specifically annotated as flowering (e.g. “flowering”, “flr”, “in flower”) and occurred in North America from 1900-2023. These records were downloaded from the Consortium of Midwest Herbaria, a database in the SEINet Portal Network (https://midwestherbaria.org/portal/index.php) and species’ names were updated in accordance with Kew World of Plants accepted species’ names. Records were then further filtered to exclude duplicate records as well as those records that were missing data in the following categories: species name, occurrence coordinates, and date collected (day, month, and year). Species with less than 50 individual specimens were also removed from the study. The final dataset consisted of 14,343 herbarium records for 52 species.

Because we only used herbarium records annotated as flowering in our main analyses, many other digitized herbarium specimens were excluded from our study. To cross-check the accuracy of digital annotations and to evaluate whether annotated specimens were representative of all available specimens, for one target species (*Viola pubescens*) we manually assessed the flowering status of all imaged herbarium specimens—a dataset that included both specimens annotated as flowering and specimens lacking this metadata. This assessment confirmed that flowering data from annotated specimens was strongly representative of flowering data from all specimens. We found that both the annotated specimen dataset (n = 220) and the expanded/manual dataset (n = 431) showed a significant advance in flowering day over time for this species. The inferred flowering shift in our manually collected flowering dataset (slope = – 0.316 days/year, p = 5.68e-17, Figure S1) was somewhat stronger than the shift detected in the annotated specimens only (slope = –0.196, p = 9.74e-5)—the data used in our main analysis. This indicates that our main analyses draw on estimates of flowering time shifts that are qualitatively representative of, and possibly more conservative than, estimates from all herbarium records.

### Historical climate data

We obtained monthly mean temperature and monthly total precipitation data for the past 124 years (1900-2023) from the Parameter-elevation Regressions on Independent Slopes Model (PRISM) climate dataset at a 30-arcsecond resolution (PRISM Group, Oregon State University). Because PRISM data exists only for the contiguous United States, occurrences outside of this area were excluded from this analysis, which resulted in a total of 13,857 records for these analyses. For each remaining herbarium record, we calculated the mean temperature and the total precipitation for the 12-month period prior to the flowering occurrence/collection date. For example, for a flowering occurrence on March 15, 1950, we calculated the mean temperature (°C) and total precipitation (mm) from March 1949 - February 1950. We used the entire preceding year of environmental data because most *Viola* species within this study are perennial (n=48) compared to annual (n=3) and therefore experience all seasons in the preceding year.

### Species’ trait data

Species trait data for mating system (CHCL versus CH), origin status (native versus introduced), life history (annual versus perennial), and flower color was collected from The Flora of North America (eFloras.org) and from a recent taxonomic treatment of violets in northeastern US (Ballard et al., 2023). To calculate species’ range sizes, we used a larger occurrence dataset that provided higher resolution to describe species ranges. For this, we downloaded specimen records with coordinates for all North American *Viola* species from Global Biodiversity Information Facility (n=60,364) (GBIF.org). After removing duplicate records, the final occurrence dataset consisted of 40,158 records for our 52 study species. Using the ‘sf’ package in R, each record within each species was given a 30km buffer; individual buffers were joined together when overlapping and then aggregated to create polygons that represent an estimate of a species’ geographic distribution. This is a relatively conservative estimate of range, as it restricts the inference to known/observed locations (and a buffer around these). We then categorized species into three continent-wide geographical ‘region’ categories—east, west, and both— depending upon whether their GBIF occurrence data were observed east or west of the 100^th^ meridian West, which is a bioclimatic boundary that divides the humid eastern and arid western US (Seager et al., 2018; Ramirez-Parada et al., 2025). Greater than 85% of occurrences must be on either side of this boundary for a species to be categorized as “eastern US” or “western US”; the remaining species (n=7), present in both regions, were categorized as “both”. Finally, using our species’ estimates of geographic range, we calculated percent range overlap for each species pair using the smaller range as the denominator. All trait data for each species is provided in the Supplementary material (Table S1).

### Statistical analyses

#### How has flowering phenology and climate shifted since 1900 in North American violets?

To quantify the shift in flowering phenology over the past 124 years in each of our 52 *Viola* species individually, we used a linear regression to model specimen flowering Julian date (i.e. day of year) as a function of year (1900-2023) for that species’ flowering records (Figure S2). We then used the slope of this regression to represent a rate of change in flowering day over time—expressed as the number of days flowering advanced or delayed per year—for each species. The collection date of the herbarium specimen is used as the flowering date, since all specimens in this analysis are flowering. For each species we also measured the correlation between latitude and collection year using a Spearman’s rank correlation to evaluate evidence for historical changes in where species were collected over our 124 time period (see Supplementary material; Table S2). To exclude the potential confounding effects that might result from this, we ran our analysis again after excluding two species with high correlation coefficients (r>0.4), and found that our results were qualitatively indistinguishable with or without these species (Supplementary material; Figure S3).

We also quantified the change in climate variables across the total range of North American violets from 1900 to the current day, using all herbarium records whose geolocations had matching PRISM data. We calculated the mean annual temperature and total annual precipitation for two periods of time: 1900-1910 and 2013-2023, then mapped the difference in each climate variable between the two periods, to capture how they had changed at the known occurrence locations in our study.

#### Are changes in two major environmental factors associated with flowering phenology? Are these consistent predictors of advanced or delayed flowering?

To test whether flowering phenology could be explained by environmental factors, we analyzed the effect of collection year, latitude, temperature, and precipitation on specimen flowering day using a nested linear regression. For each species, we ran two linear models, a simple model (S) and a more complex environmental model (E) and used an ANOVA to determine which model best represented the data. The simple model analyzed flowering day as a function of collection year and latitude. The environmental model assessed the same relationship with the addition of two environmental factors as predictor variables: mean temperature (°C) and total precipitation (mm) from the previous 12-month period. These two environmental factors were chosen as they capture key elements of climate change across North America since 1900 (see above, and Results).

We used the simple model to evaluate variation in flowering day due to year and latitude alone. Latitude is expected to affect the timing of flowering, with individuals at lower latitudes flowering earlier than higher latitudes due to earlier springs. The environmental analysis was used to assess whether environmental variables—temperature and precipitation—could explain variation in flowering day within each species, beyond the effect of latitude. We then summarized whether specific variables were consistently associated with advancing or delaying flowering across species.

#### Is variation in phenological shifts among species explained by species’ traits?

To evaluate whether species’ traits could explain variation in the overall rate at which species have advanced or delayed flowering, we used an ANCOVA to test the association between variation in species’ traits and their rate of phenological change (days/year); the latter estimate corresponds to the slopes produced in the first analysis (Fig. 1). This analysis included the following traits as predictor variables: range size (km^2^), flower color (white, yellow, violet), region (east, west, both), origin status (native, introduced), mating system (CH, CHCL), and life history (annual, perennial). We ran these models on two sets of species: first, on all species regardless of whether they were inferred to have significant shift in flowering phenology; and, second, on only the subset of species that had significant shifts in flowering phenology. Post hoc comparisons using the Tukey test were performed for species traits with a significant effect. Results for the analyses that include only significantly shifted species are reported in supplementary material (Figure S4, Table S3, Table S4), and qualitatively agree with results of analyses that include all species.

**Figure 1.**
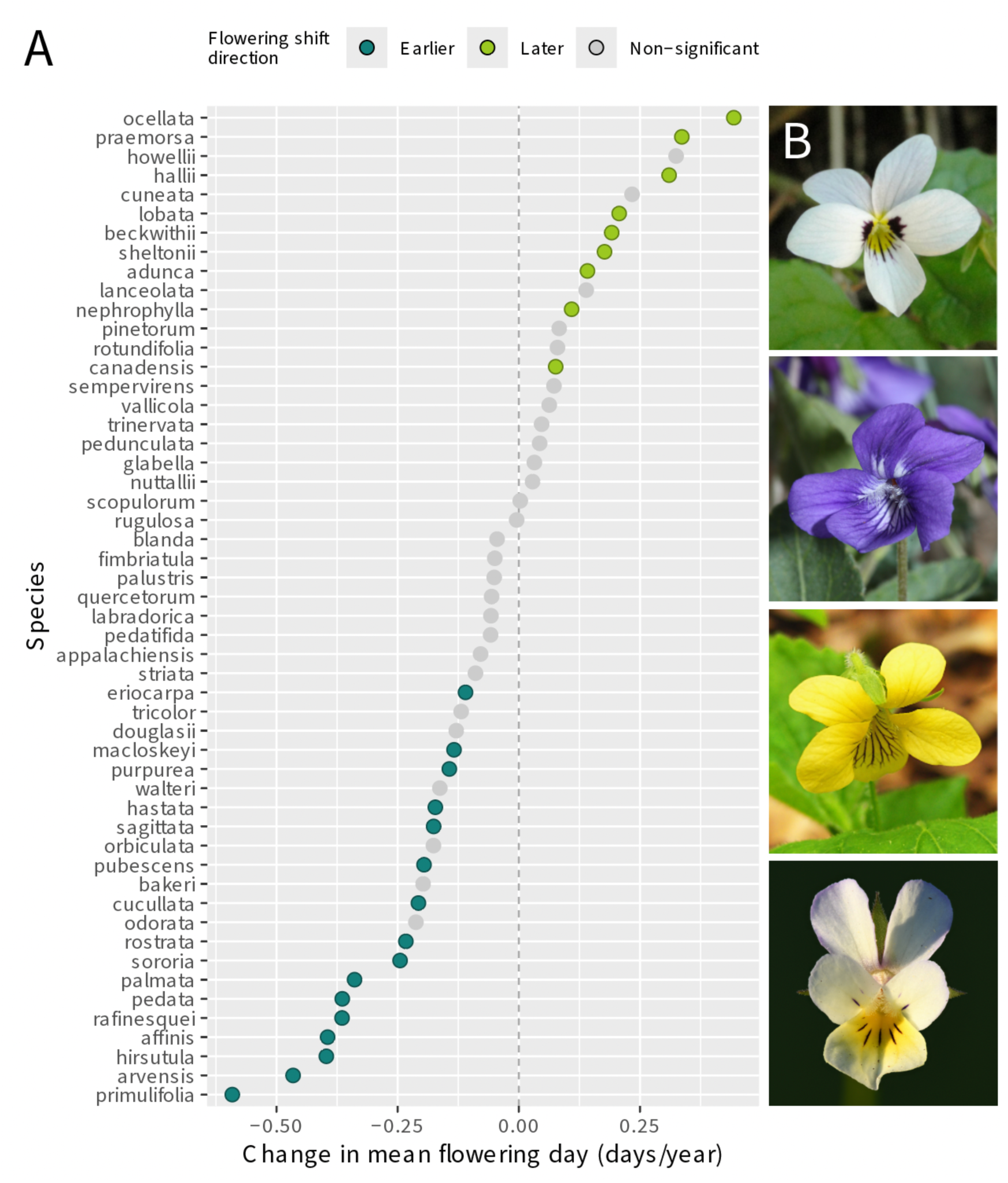
A) The rate of change in mean flowering day (days/year) for each species, based on samples from 1900-2024. The slope of the linear regression (flowering day ∼ year) is shown for each species, with negative values indicating advance in mean flowering day and positive values indicating delay in mean flowering day. Colored = significant (p<0.05), grey = not significant (p>0.05). B) Flowers of two species that delayed flowering – *V. ocellata and V. adunca* – and two species that advanced flowering – *V. pubescens and V. arvensis.* Photo attributions from top to bottom: *Viola ocellata*, Miguel Vieira, CC BY 2.0; *Viola adunca*: Walter Siegmund, CC BY-SA 3.0; *Viola pubescens*: Nicholas A. Tonelli CC BY 2.0; *Viola arvensis*: Ivar Leidus, CC BY-SA 4.0. Accessed via Wikimedia Commons.

#### Have shifts in flowering phenology changed patterns of co-flowering among sympatric species?

We investigated how species’ changes in flowering time from historical to current day have affected patterns of pairwise co-flowering, using a subset of species pairs that had sufficient data on range and flowering date. To do so, we defined the peak flowering window (middle 80% of data) for a historical (1900-1923) and current day (2000-2023) time period for each species. The middle 80% of data was used as it captures when a species is most abundantly flowering and most likely to be reproductively interacting with other species. Only species with geographic range data and >20 flowering records for both time periods were included, resulting in sixteen species and therefore 120 species pairs for these analyses. For each species pair, to summarize how their potential for phenological overlap has changed since 1900, we calculated the length of the co-flowering window (i.e. the overlap in the two species peak flowering windows, in days) during each of the historical (1900-1923) and current (2000-2023) time periods. The change in co-flowering between these two time periods was then calculated as the difference in length of the historical and current co-flowering windows (in days), which is shown in main results. Because the length of a flowering window can be very different between species, we also visualized the change in co-flowering as the difference in the proportion of species A’s flowering window that overlapped with species B’s window from 2000-2023 to 1900-2023, where species A is the focal species (See supplementary text; Figure S6).

To address whether pairwise co-flowering distributions have significantly changed over time, for each species pair we tested whether co-flowering windows have become significantly more narrow or broad between the historical and current time periods. Specifically, for each species pair, flowering occurrence records of both species that fell within their co-flowering window were combined to form a collective co-flowering distribution for each of the historical and contemporary periods (as illustrated in Figure S5). We then conducted a Levene’s test to determine whether the variance differed significantly between the historical and current co-flowering distributions, for each species pair. Statistical tests were Bonferroni-corrected to account for multiple comparisons.

## Results

### How has flowering phenology and climate shifted since 1900 in North American violets?

Flowering phenology has shifted significantly for 25 of the 52 species of North American *Viola* in our study (Fig. 1). Sixteen species have shifted flowering earlier, with estimated advances in flowering ranging from 0.11-0.59 days/year. This equates to average flowering day advancing by between ∼14 and 73 days since 1900. Nine species have shifted flowering later, with estimated delays in flowering ranging from 0.08-0.44 days/year, which equates to flowering of between ∼10 and 55 days later since 1900 (Fig. 1). Across this same 124 year period, we found substantial shifts in environmental variables across the species’ ranges, when comparing 10 year temperature and precipitation averages between historical (1900-1910) and recent (2013-2023) time windows (Fig 2). Mean annual temperature has increased for almost every occurrence location, ranging from -0.8 to +3.2 °C, with a median temperature change of +0.87°C. In comparison, total annual precipitation has changed variably across North America during the same 124 year period, increasing by up to ∼50cm in some regions (notably in the eastern US) and decreasing by ∼130cm per year elsewhere, especially in parts of California (Fig. 2). This represents a 1.6-fold increase and >2-fold decrease in precipitation in the most extreme cases. *Are changes in major environmental factors associated with flowering phenology? Are these consistent predictors of advanced or delayed flowering?*

**Figure 2.**
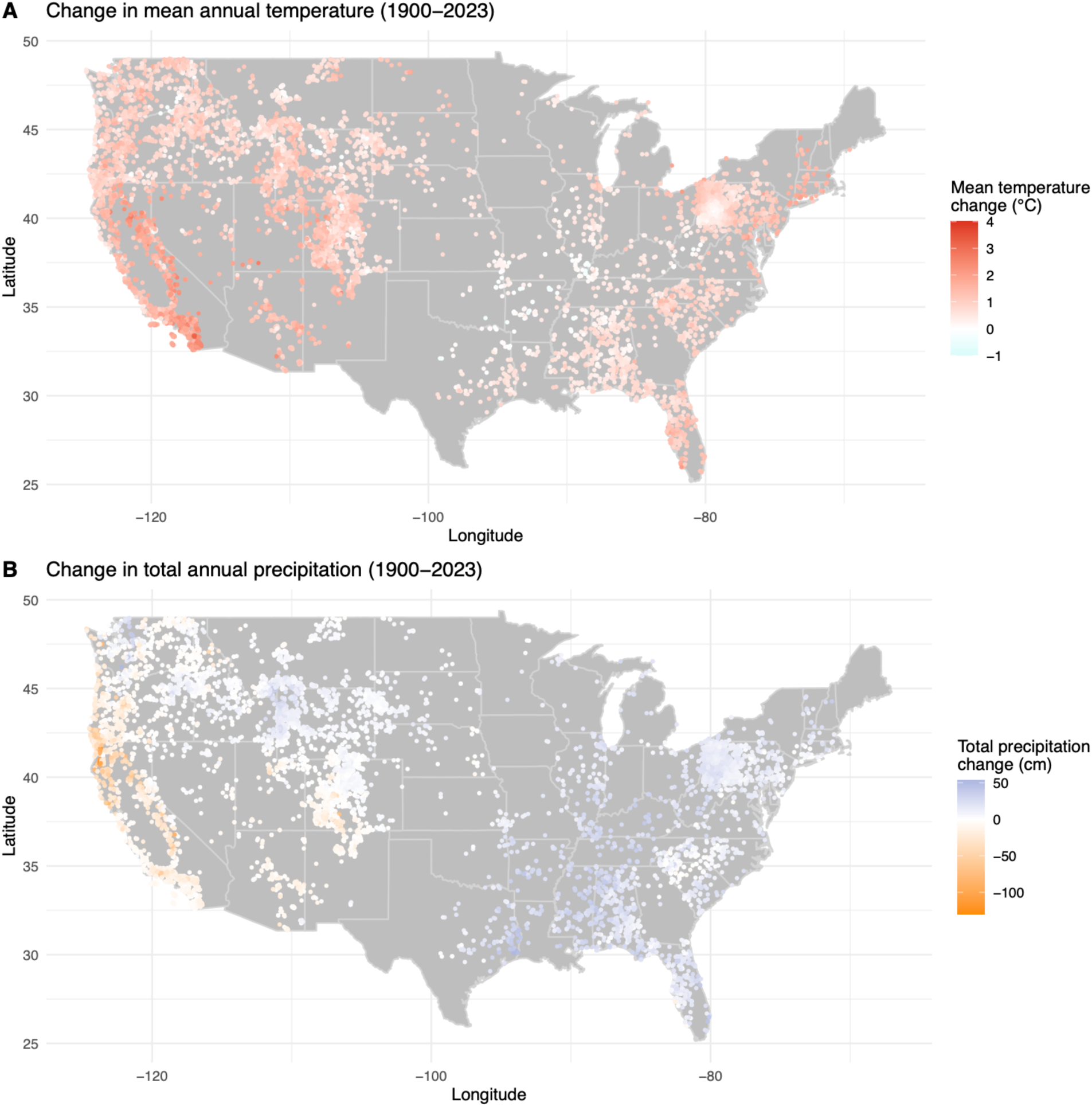
Map of all *Viola* occurrence records used in analysis of environmental associations with phenology shifts, colored by change in A) mean annual temperature (°C) and B) total annual precipitation (cm) between the periods 1900-1910 and 2013-2023.

An environmental model (with year, latitude, temperature, and precipitation) fit the data significantly better than a simple model (with only year and latitude) for the majority of species in our dataset (Table S5). Of 52 species, the environmental model was significant for 46 species and the best fit model for 40 species (Fig 3). In comparison, while the simple model was significant for 36 species, it was the best fit model for only five species. (For six species, neither model was significant.) Similarly, when considering only those 25 species with significant shifts in flowering phenology (Fig 1), most (n=19) were best explained by the environmental model (Fig 3) including both earlier (n=11) and later (n=8) flowering species. In comparison, five species—all of which significantly advanced flowering—were best explained by the simple model (Fig 3).

**Figure 3.**
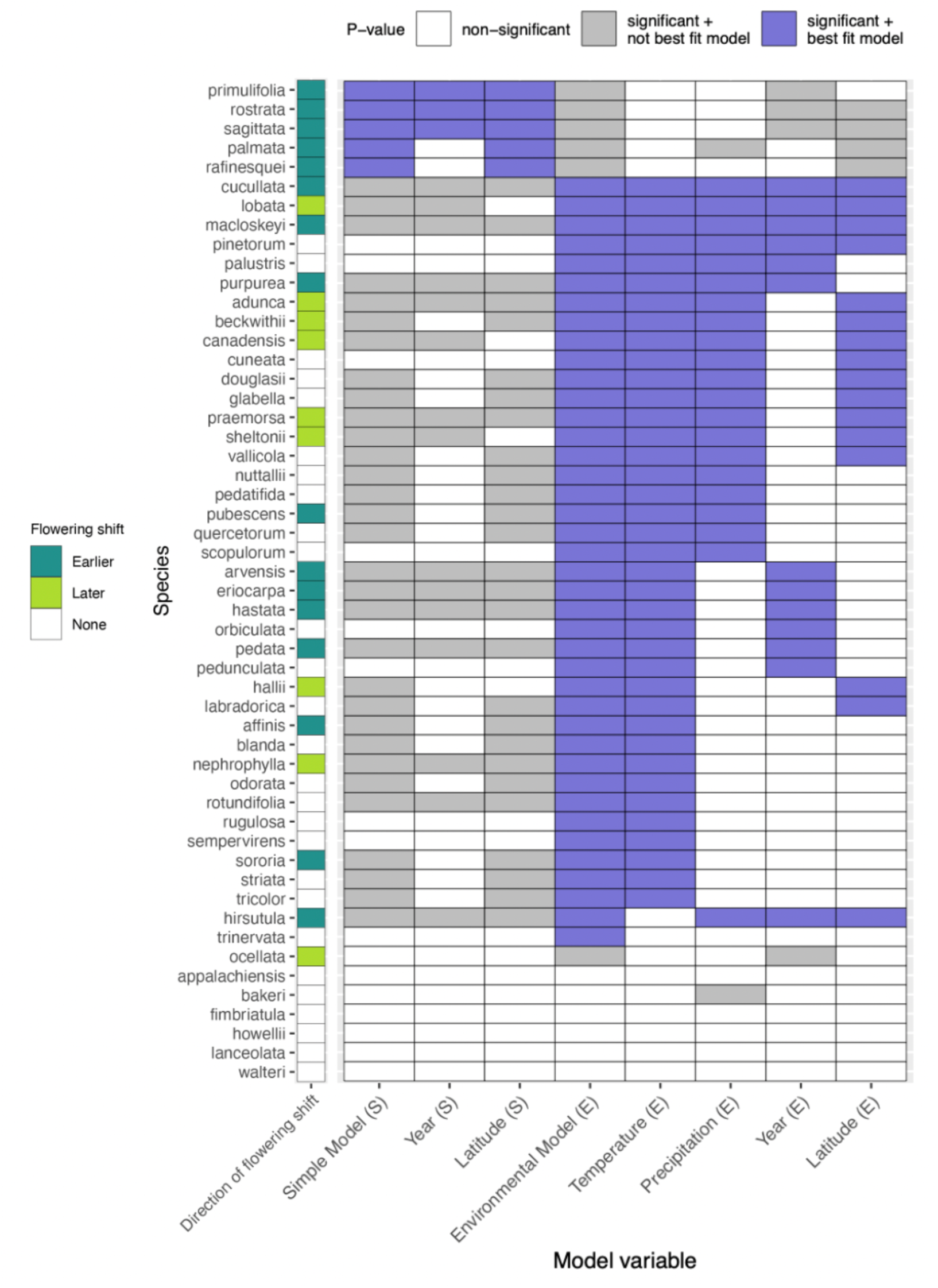
Results of nested linear models, comparing the simple model (S) and the more complex environmental model (E). Model S: flowering day ∼ year + latitude. Model E: flowering day ∼ temperature + precipitation + year + latitude. The furthest left column summarizes flowering shift for each species (dark green = earlier flowering, light green = later flowering; see Figure 1). The other columns represent the significance of the overall models (B or E) or model variables (purple = significant + best fit model, grey = significant + not best fit model, white = non-significant).

Considered across models and species, temperature was the predictor variable that is most explanatory of flowering day. For all species for which the environmental model was best fit (n=40), most (n=38) had a significant temperature effect (Fig 3). In half of these cases (n=20), precipitation was also significant in the same model (Fig 3). Temperature effects were also observed in conjunction with year (n=6) and latitude (n=2) effects, or with more than one additional factor (n=16). In 10 cases, temperature alone was explanatory. After temperature, precipitation was the second most explanatory variable (Fig 3). Of 21 cases with significant precipitation terms, almost all were observed in conjunction with at least one (n=5), two (n=12), or three (n=4) additional significant main effects (Fig 3).

These patterns of explanatory environmental effects were similar when considering only species whose flowering date was significantly shifted over time (n=19) (Fig 3). Of the 11 species with significantly earlier flowering, 10 had a significant temperature effect; four of these cases were in conjunction with a precipitation effect, and four were with a year effect (Fig 3). Only one species showed a significant effect of precipitation without temperature. Temperature was also significant for all eight species with delayed flowering; in most cases (n=6) this effect occurred in conjunction with precipitation and at least one other main effect (latitude and/or year).

### Is variation in phenological shifts among species explained by species’ traits?

Of six assessed species traits, the only significant predictor of mean flowering shifts was region (p=0.0003; Table 1). This effect was observed regardless of whether our analysis included all species, or just the subset of species that had significant shifts in flowering phenology over time (Table S3). This effect is explained by a significant difference in the rate of flowering shift specifically between eastern and western species (p = 0.0002; Table 2); species found in both regions did not differ from either ‘east’ or ‘west’ species (Table 2). Eastern and western species differ in their average rate of flowering shift by 0.264 days/year, due to the combined effect of eastern species largely advancing flowering and western species predominantly delaying flowering. On average, eastern species advanced flowering by 0.190 days/year (∼24 days since 1900) and western US species delayed flowering by 0.075 days/year (∼9 days since 1900). No other species trait (of 6 total) evaluated in our ANCOVA analysis had a significant effect on mean flowering shift (Table 1, Table S6); a subset of representative traits is shown in Figure 4 and all traits are shown in Figure S7.

**Figure 4.**
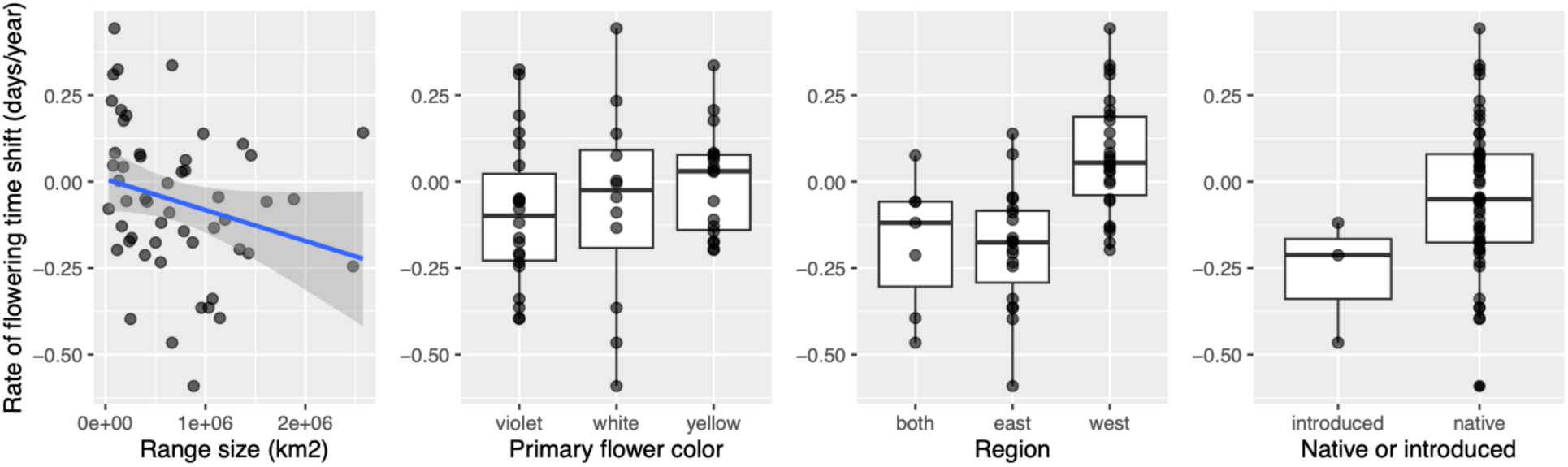
The rate of flowering phenology shifts (days/year) analyzed by A) range size (p = 0.26), B) flower color (p = 0.65) C) region (p = 0.0003), and D) origin status (p = 0.64). Each point represents a species’ rate of flowering change (from Figure 1).

**Table 1.** ANCOVA results for the effect of species’ traits on rate of flowering time shift for all species. Model: rate of flowering shift ∼ range size + flower color + region (east/west/both) + origin (native/introduced) + mating system + life history (annual/perennial). * = significant.

|  | *Sum Sq* | *Df* | *F value* | *p-value* |
| --- | --- | --- | --- | --- |
| *Range size* | 0.04078 | 1 | 1.2972 | 0.2610 |
| *Flower color* | 0.02731 | 2 | 0.4343 | 0.6505 |
| *Region* | 0.60969 | 2 | 9.6966 | 0.0003 * |
| *Origin* | 0.00690 | 1 | 0.2195 | 0.6418 |
| *Mating system* | 0.00511 | 1 | 0.1627 | 0.6887 |
| *Life history* | 0.02378 | 1 | 0.7564 | 0.3893 |

**Table 2.** Tukey posthoc contrast for species trait ‘Region’. The estimate shows the difference in flowering shift rate (days/year) between species in the three region categories (east/west/both).

|  | *Estimate* | *Std. Error* | *t value* | *p-value* |
| --- | --- | --- | --- | --- |
| *East - Both* | -0.07797 | 0.09942 | -0.784 | 0.7074 |
| *West - Both* | 0.18571 | 0.10421 | 1.782 | 0.1806 |
| *West - East* | 0.26368 | 0.05993 | 4.400 | 0.0002 * |

### Have shifts in flowering phenology changed patterns of co-flowering among sympatric species?

Sympatry is common among the 109 species pairs for which we had sufficient flowering records and range size data (see Methods): 76 species pairs have 20% or more range overlap, and 33 of these have >40% range overlap. Accordingly, shifts in co-flowering among these species includes many pairs that could directly interact. Of all pairs, approximately one third have increased the number of co-flowering days (n=35) in the last 124 years while two thirds have experienced a decrease in the number of co-flowering days (n=73), including a complete loss of co-flowering for 15 species pairs (Table S7). Figure 5 shows these data in terms of the absolute difference (present – past, in days) between co-flowering windows, against the estimated range overlap between each species pair; these changes in co-flowering overlap range from a decrease of 40 days to an increase of 59 days. Our tests evaluating whether these represent significant changes between historical (1900-1923) and current (2000-2023) co-flowering found that historical and current co-flowering variances differed significantly for 48 species pairs (denoted by colors in Figure 5A), after correcting for multiple comparisons. About a third of these (n=17) involve a significant increase in current flowering overlap, almost all of which are in species pairs with a >40% range overlap (n=15) (Figure 5). The other two thirds of species pairs (n=31) decreased in co-flowering; the majority (n=21) of these have low range overlap, but range overlap is >40% for ten pairs (Figure 5A). To further understand whether specific co-flowering changes are more prominent in different locations, we also identified the geographic context for these co-flowering shifts. We found that increased co-flowering is occurring primarily in the eastern US and decreased co-flowering is occurring primarily in western US (Figure 5B). Widespread species are also more likely to influence patterns of range overlap, and potentially co-flowering, between species; Figure 5B also highlights two widespread species to identify their contribution to overall co-flowering: *Viola sororia* in eastern US and *Viola adunca* in western US.

**Figure 5.**
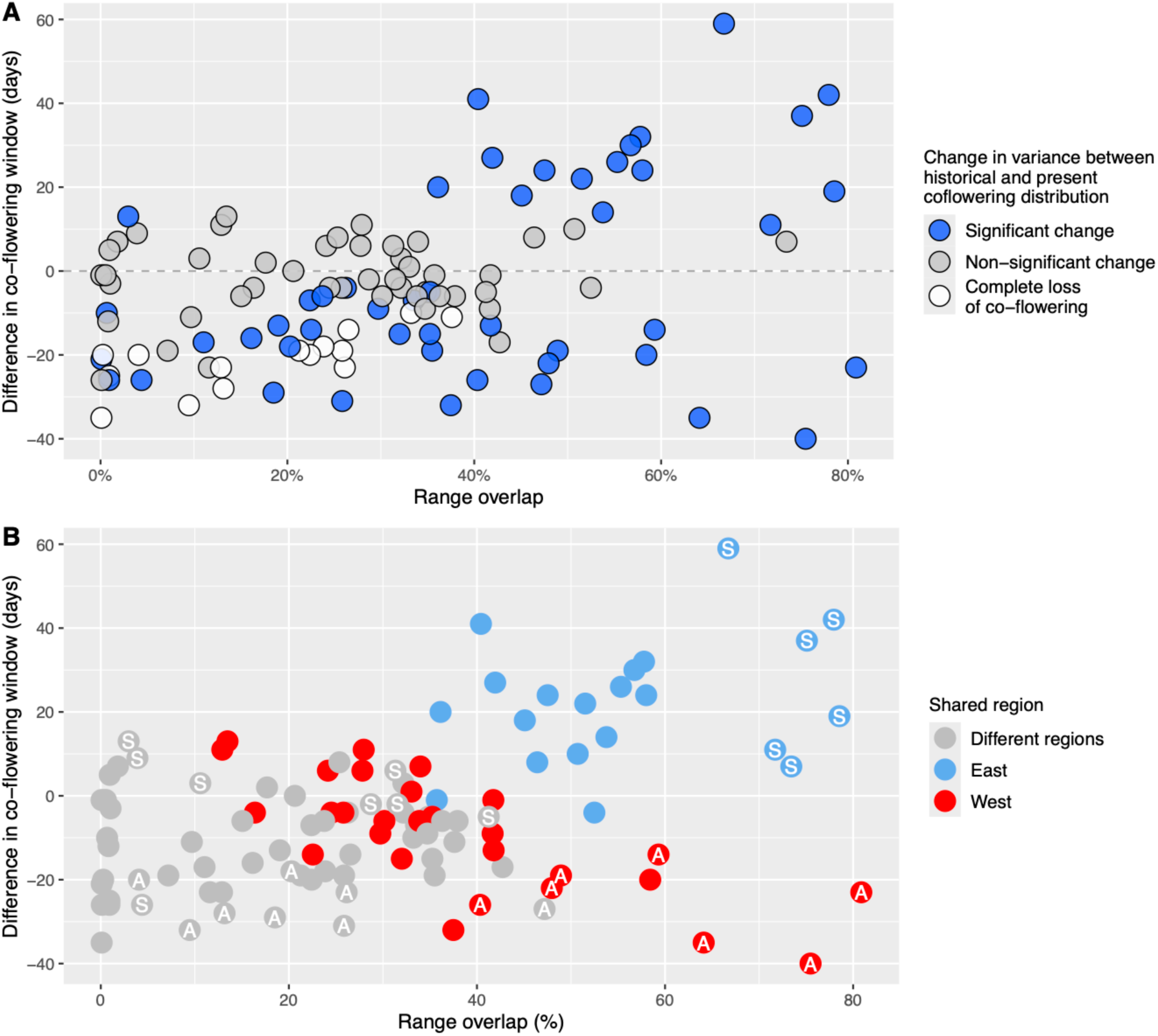
The relationship between percent range overlap and the change in flowering overlap (number of days that pairs are co-flowering) from 1900-1924 (past) to 2000-2024 (present) for all species pairs with >0% range overlap. (Pairs with 0% overlap (n=11) were omitted for visual clarity.) Each point is a species pair. A) Species pairs for which the variance of collective co-flowering significantly differed between past and present time periods are colored blue; non-significant comparisons are grey. Species pairs that historically co-flowered but no longer co-flower are white. B) Species pairs colored by the shared region among both species in a pair. The subset of species pairs that include one of two widespread species, in the east (*Viola sororia*) and the west (*Viola adunca*), are labelled with S (*Viola sororia*) or A (*Viola adunca*).

Finally, with these data we are not able to directly quantify the effect of co-flowering shifts on hybridization or gene flow, but we can examine one correlate of the potential for reproductive contact: flower color. Specifically, species that overlap geographically are more likely have pollinator movement between them, and therefore experience hybridization, when they share flower color. When we assess the distribution of shared flower color among species pairs, we find that of the pairs with large (> 40%) range overlap and increased co-flowering— those pairs most likely to have increased reproductive interaction under climate change—more than half (10/18) share the same flower color (Figure S8).

## Discussion

Using ∼14,000 flowering occurrence records for 52 species of North American *Viola*, we found that 1) variable and in some cases very large phenological shifts have occurred in violet species since 1900, 2) the environment is significantly and consistently associated with these shifts, specifically temperature and geographic region of US and 3) these shifts have resulted in substantial changes to co-flowering among species pairs, ranging from an increase of 59 days to a complete loss of co-flowering. Below we discuss these findings, in the context of understanding the ecological and physiological drivers of phenological change in this and other plant systems, and the potential evolutionary outcomes of those changes.

### Large phenological shifts accompany environmental changes over the last 124 years in North American violets

Concordant with estimates of contemporary climate change (Easterling et al., 2017; Vose et al., 2017), we confirmed that key environmental variables—temperature and precipitation— have experienced substantial and spatially variable changes across the geographical range of North American *Viola* since 1900 (Figure 2). In response to this changed climate, species in this group exhibit a striking range of individual responses within a collective pattern of altered flowering time. The magnitude of these phenological shifts are comparable to estimates of flowering shifts worldwide, which show an average advancement of 0.247 days/year and an average delay of 0.278 days/year (Williamson et al., 2025). Prior studies that span large geographic regions have found similar patterns of heterogeneity in phenological response to climate change among species (Zu et al., 2025; König et al., 2018). However, because studies that have wide geographic sampling often also have wide taxonomic representation, interpreting the drivers of this heterogeneity can be challenging, as relevant trait differences can be confounded with many other species differences. Conversely, studies focused on close relatives—which have found that congeners share similar phenological responses to climate change (Carscadden et al., 2022; Park et al., 2022)—generally sample few species per genus (as few as two) making it more difficult to discern general patterns. By assessing dozens of North American *Viola* species, we uncover a clear pattern of heterogeneity in phenological responses among many closely related species across a contiguous geographic region. This indicates that the heterogeneous phenological responses often observed across broader taxonomic scales can and do arise across relatively short evolutionary timescales (Marcussen et al., 2022), among species within the same genus. It also allowed us to identify general patterns of response to climate change, while evaluating species traits that vary at the intrageneric level as potential causes of this heterogeneity.

### The intersection of environment and geography explains broad patterns of phenological responses in Viola

Despite heterogeneity in both the direction and magnitude of flowering day shifts among species, we still detected strong, systematic effects of two factors on *Viola* phenology: continent-level geographic region, and local temperature. The latter finding is consistent with many previous studies that have found associations between flowering shifts and temperature (Menzel et al., 2006; Miller-Rushing & Primack, 2008; Austin et al., 2024). Increasing temperature is widely associated with earlier flowering, including (like *Viola*) in spring-flowering species (Menzel et al., 2006; Miller-Rushing & Primack, 2008; Yu et al., 2010; Bartlett et al., 2023), and a majority of our significantly shifted species showed advances in flowering date (Figure 1). Nonetheless, we also found that one third of *Viola* species significantly delayed flowering, despite almost all geographic ranges experiencing increased temperature since 1900 (Figure 2). Several pieces of evidence indicate that these delayed flowering responses are shaped by other environmental changes that could have independent effects on flowering timing. Precipitation in particular has been shown to have more complex effects on flowering phenology compared to temperature (Matthews & Mazer, 2016; Mazer et al., 2013). In some cases, increased precipitation has been associated with delayed (Von Holle et al., 2010) or advanced flowering (Crimmins et al., 2010; Lambert et al., 2010), and in other cases, change in precipitation has no association with flowering shifts (Sherry et al., 2007). Consistent with these observations, in *Viola* we found that precipitation was a significant predictor of flowering date for several species, but often in conjunction with other explanatory main effects, including year, latitude, and temperature.

More importantly, however, these more complex environmental effects intersect directly with our only other significant predictor of flowering shift in North American *Viola*—geographic region. We found that western US species on average delayed flowering by 0.264 days/year (∼33 days since 1900) compared to species in the eastern US, which on average advanced flowering (p = 0.0002, Table 2; Figure 4). For delaying species specifically, the combined effect of temperature and precipitation explain flowering time for two thirds of species (Figure 3). Taken together with the observation that a large fraction of the western occurrences show decreases in total annual precipitation (Fig. 2), we infer that delays in flowering are a frequent *Viola* response to the joint environmental effects of increasing temperatures and decreasing precipitation, a combination that commonly occurs in western but not eastern US.

In comparison to these factors, none of the other species-level traits that we assessed (range size, flower color, mating system, origin status, and life history) significantly explained variation in observed phenological responses. Origin status and life history traits have been associated with shifts in flowering phenology in other systems (see *Introduction*). We did not find these patterns in *Viola,* although our study included relatively few introduced (n=3) and annual (n=4) species, which might have reduced our ability to discern these effects. Reproductive traits that affect reliance on external pollinators have also been associated with shift in flowering phenology in previous studies (see *Introduction*), but we also found no differences in flowering shifts between CHCL and CH species. Accordingly, variation in intrinsic species traits provides a poor general explanation for the diverse phenological responses we observe among these 52 species.

Instead, our findings indicate that *Viola* species’ phenological shifts over the past 124 years are most strongly influenced by systematic responses to several critical environmental factors—especially temperature but also precipitation—in conjunction with where species are found—that is, by their geographic region. The latter is important because regional differences within North America determine how these critical elements of the environment have been altered by recent climate change (Fig 2).

Finally, in addition to these clear explanatory effects, our models suggest that other climate variables likely also contribute to some unexplained phenological heterogeneity in our dataset. For instance, in addition to temperature and precipitation, 11 advancing and two delaying species also showed a significant effect of year (Figure 3), which most likely reflects flowering phenology shifts arising from changes in other unmeasured environmental factors that are highly correlated with recent time. These could include changes in environmental variability (Pendergrass et al., 2017), frequency of extreme events (Vázquez et al., 2017), or unfulfilled chilling requirements (Yu et al., 2010), among other more complex factors. Nonetheless, on the background of these effects, our findings underscore that the interaction between precipitation and temperature can be important for flowering phenology (Matthews & Mazer, 2016), and—in the context of regional variation in these factors—contribute to both advanced and delayed flowering responses to climate change.

### Significant increases and decreases in co-flowering accompany changing climate

Climate-driven phenological shifts have been shown to alter patterns of co-flowering in many different plant communities (CaraDonna et al., 2014; Forrest et al., 2010; Austin et al., 2024; Pareja-Bonilla et al., 2025), but the specific consequences of these shifts for altering reproductive contact between congeneric community members is much less clear. Here our results indicate that species’ phenological shifts in response to climate change have had significant effects on patterns of species co-flowering, with potentially large effects on interactions among co-occurring *Viola*.

First, and consistent with other observations of co-flowering shifts within communities (Austin et al., 2024; CaraDonna et al., 2014; Fisogni et al., 2022), we find flowering shifts have a spectrum of effects on co-flowering among *Viola* species, including some pairs significantly increasing and others decreasing co-flowering. However, when region is simultaneously considered, we observe a strong geographic signal in the direction of these changes, where increases in co-flowering are common among pairs in eastern US whereas species pairs in western US tend to decrease co-flowering (Figure 5B). This is in part driven by widespread species (e.g. *Viola sororia*, *Viola adunca*) whose flowering shift impacts all species that co-occur throughout their range. Regardless, the distinctive influence of geographical region effects is again apparent in our co-flowering data, just as it was in shaping individual species phenological responses.

Second, our finding that more than half of the species pairs with large range overlap (> 40%) and increased co-flowering also share the same flower color indicates these shifts could have large impacts on pollinator mediated interactions among *Viola* species. Ultimately, the evolutionary consequences of shifts in co-flowering depend upon the biological significance of co-flowering itself. For instance, reduced co-flowering might be beneficial, if co-flowering otherwise leads to deleterious hybridization (Chunco, 2014). Conversely, increased co-flowering might have beneficial effects, such as increasing the attraction of pollinators (Sargent & Ackerly, 2008). However, several features of *Viola* suggest that increased co-flowering is more likely to have deleterious than beneficial effects. For instance, *Viola* species are pollinated by generalist insects, making pollinator limitation (and therefore the positive, facilitating effects of co-flowering) less likely. Moreover, reproductive barriers among *Viola* species are already known to be fragile, with 80 different hybrid types historically documented in the wild (Brainerd, 1904; Ballard et al., 2023). Climate-driven increases in reproductive overlap between species risks even greater disruption of species boundaries in the future. Overall, while the specific outcomes of *Viola*’s large and varied shifts in flowering and co-flowering phenology will likely differ between individual species pairs, we have shown here that the opportunity for pollinator-mediated reproductive interactions has changed over time, and that a substantial fraction of these species pairs could experience increased reproductive interaction under climate change. These findings underscore the potential for climate-driven phenological shifts to have large and long-term evolutionary consequences in groups of closely related, geographically contiguous species like North American *Viola*.

