## Supplemental materials for "Large phenological advances and delays over 124 years of climate change alter co-flowering among North American *Viola*"

### Supplemental methods

*Correlations among latitude and year due to herbarium collecting patterns*

For each species individually, we assessed whether observed patterns of flowering time shift could be spuriously influenced by geographical changes in patterns of specimen collecting over time. To do so, we analyzed correlations among latitude and year collected for each species, using Spearman's rank correlation method (Table S2). Since latitudes differ in flowering times (e.g. lower latitudes flower earlier), we tested whether latitude and year were strongly correlated, and therefore whether detected shifts in flowering over time could simply reflect a change in geographical collecting locations over time, for each species. Specifically, if species were inferred to advance flowering, we assessed for a strong negative correlation (correlation coefficient larger than -0.4) between year and latitude, and for delaying species, we assessed for a strong positive correlation (correlation coefficient >0.4) between year and latitude.

For 50 of 52 species, we did not find evidence for a strong correlation between year and latitude. These factors were significantly and strongly correlated for two species only – *Viola pedata* and *Viola hirsutula* – both of which were advancing species (Table S2, Figure S3).

*Calculating change in co-flowering as a proportion of total flowering window*

We primarily characterized the change in co-flowering between contemporary and historical time periods as the difference in length of the historical and current co-flowering windows (see main text). To also investigate change in co-flowering as a proportion of overall flowering window length, we calculated the change in flowering overlap using the following equation:

$$change\ in\ flowering\ overlap = \frac{Overlap\ between\ A\ and\ B\ (2000-2023)}{Flowering\ period\ of\ A\ (2000-2023)} - \frac{Overlap\ between\ A\ and\ B\ (1900-1923)}{Flowering\ period\ of\ A\ (1900-1923)}$$

where species A is the focal species. The resulting values range from -1 to 1 where positive values indicate an increase in flowering overlap and negative values indicate a decrease in flowering overlap between historic (1900-1923) to current (2000-2023) time periods (Figure S5).

### Supplemental Tables and Figures

Table S1. Species trait data: range size ($km^2$), flower color (white, yellow, violet), region (east, west, both), origin status (native, introduced), mating system (CH, CHCL), and life history (annual, perennial)

| Species | Mating system | Origin | Life history | Region | Primary petal color | Petal length (mm) | Range size ($km^2$) | Source |
|---|---|---|---|---|---|---|---|---|
| *adunca* | CHCL | native | perennial | west | violet | 17 | 2575992 | eFloras |
| *affinis* | CHCL | native | perennial | both | violet | 22 | 1144273 | eFloras |
| *appalachiensis* | CHCL | native | perennial | east | violet | 18 | 31811 | eFloras |
| *arvensis* | CH | introduced | annual | both | white | 15 | 665836 | eFloras |
| *bakeri* | CHCL | native | perennial | west | yellow | 14 | 116969 | eFloras |
| *beckwithii* | CH | native | perennial | west | violet | 22 | 209725 | eFloras |
| *blanda* | CHCL | native | perennial | east | white | 10 | 1128969 | eFloras |
| *canadensis* | CHCL | native | perennial | both | white | 20 | 1455865 | eFloras |
| *cucullata* | CHCL | native | perennial | east | violet | 13 | 1430771 | eFloras |
| *cuneata* | CHCL | native | perennial | west | white | 14 | 62731 | eFloras |
| *douglasii* | CH | native | perennial | west | yellow | 21 | 163285 | eFloras |
| *eriocarpa* | CHCL | native | perennial | east | yellow | 18 | 1194844 | eFloras |
| *fimbriatula* | CHCL | native | perennial | east | violet | 15 | 398312 | eFloras |
| *glabella* | CHCL | native | perennial | west | yellow | 18 | 796464 | eFloras |
| *hallii* | CH | native | perennial | west | violet | 18 | 78558 | eFloras |
| *hastata* | CHCL | native | perennial | east | yellow | 12 | 238824 | eFloras |
| *hirsutula* | CHCL | native | perennial | east | violet | 17 | 251103 | eFloras |
| *howellii* | CHCL | native | perennial | west | violet | 23 | 124006 | eFloras |
| *labradorica* | CHCL | native | perennial | both | violet | 16 | 1611443 | eFloras |
| *lanceolata* | CHCL | native | perennial | east | white | 12 | 978456 | eFloras |
| *lobata* | CHCL | native | perennial | west | yellow | 19 | 155466 | eFloras |
| *macloskeyi* | CHCL | native | perennial | west | white | 12 | 1085548 | eFloras |
| *nephrophylla* | CHCL | native | perennial | west | violet | 28 | 1375651 | eFloras |
| *nuttallii* | CHCL | native | perennial | west | yellow | 13 | 764328 | eFloras |
| *ocellata* | CHCL | native | perennial | west | white | 15 | 89510 | eFloras |
| *odorata* | CHCL | introduced | perennial | both | violet | 22 | 394668 | eFloras |
| *orbiculata* | CHCL | native | perennial | west | yellow | 17 | 501501 | Flora of North America |
| *palmata* | CHCL | native | perennial | east | violet | 25 | 1068689 | eFloras |
| *palustris* | CHCL | native | perennial | west | violet | 16 | 1883614 | eFloras |
| *pedata* | CH | native | perennial | east | violet | 24 | 1031430 | eFloras |
| *pedatifida* | CHCL | native | perennial | both | violet | 25 | 416560 | eFloras |

| | | | | | | | | |
|---|---|---|---|---|---|---|---|---|
| *pedunculata* | CH | native | perennial | west | yellow | 20 | 177736 | eFloras |
| *pinetorum* | CHCL | native | perennial | west | yellow | 12 | 96120 | eFloras |
| *praemorsa* | CHCL | native | perennial | west | yellow | 15 | 666233 | eFloras |
| *primulifolia* | CHCL | native | perennial | east | white | 16 | 881819 | eFloras |
| *pubescens* | CHCL | native | perennial | east | yellow | 18 | 1342280 | eFloras |
| *purpurea* | CHCL | native | perennial | west | yellow | 14 | 783578 | eFloras |
| *quercetorum* | CHCL | native | perennial | west | yellow | 16 | 210308 | eFloras |
| *rafinesquei* | CHCL | native | annual | east | white | 10 | 959503 | eFloras |
| *rostrata* | CHCL | native | perennial | east | violet | 20 | 551700 | eFloras |
| *rotundifolia* | CHCL | native | perennial | east | yellow | 11 | 344561 | eFloras |
| *rugulosa* | CHCL | native | perennial | west | white | 20 | 621667 | eFloras |
| *sagittata* | CHCL | native | perennial | east | violet | 15 | 872810 | eFloras |
| *scopulorum* | CHCL | native | perennial | west | white | 13 | 134911 | eFloras |
| *sempervirens* | CHCL | native | perennial | west | yellow | 17 | 350594 | eFloras |
| *sheltonii* | CHCL | native | perennial | west | yellow | 18 | 181570 | eFloras |
| *sororia* | CHCL | native | perennial | east | violet | 25 | 2475283 | Flora of North America |
| *striata* | CHCL | native | perennial | east | white | 18 | 640773 | eFloras |
| *tricolor* | CH | introduced | annual | both | violet | 15 | 557852 | eFloras |
| *trinervata* | CH | native | perennial | west | violet | 15 | 77824 | eFloras |
| *vallicola* | CHCL | native | perennial | west | yellow | 14 | 801367 | eFloras |
| *walteri* | CHCL | native | perennial | east | violet | 18 | 262594 | eFloras |

Table S2. Correlation (spearman's rho) and p-value among collection year and for each species. Species that were excluded due to potential collection bias are bolded (Figure S3). Asterisks indicate a statistically significant change in flowering time since 1900.

| Species | Estimate | P-value | Direction of Flowering Shift |
|---|---|---|---|
| *adunca* | -0.115 | 4.73E-07 | Delaying * |
| *affinis* | -0.343 | 4.25E-07 | Advancing * |
| *appalachiensis* | 0.284 | 0.023 | Advancing |
| *arvensis* | -0.277 | 0.007 | Advancing * |
| *bakeri* | -0.368 | 0.006 | Advancing |
| *beckwithii* | -0.109 | 0.275 | Delaying * |
| *blanda* | -0.061 | 0.519 | Advancing |
| *canadensis* | 0.119 | 4.17E-05 | Delaying * |
| *cucullata* | -0.037 | 0.461 | Advancing * |
| *cuneata* | -0.299 | 0.004 | Delaying |
| *douglasii* | 0.163 | 0.047 | Advancing |
| *eriocarpa* | -0.118 | 0.031 | Advancing * |
| *fimbriatula* | 0.053 | 0.656 | Advancing |
| *glabella* | -0.025 | 0.568 | Delaying |
| *hallii* | -0.402 | 0.001 | Delaying * |
| *hastata* | -0.147 | 0.087 | Advancing * |
| ***hirsutula*** | **-0.465** | **5.72E-05** | Advancing * |
| *howellii* | -0.324 | 0.018 | Delaying |
| *labradorica* | 0.100 | 0.073 | Advancing |
| *lanceolata* | 0.132 | 0.100 | Delaying |
| *lobata* | -0.386 | 1.35E-05 | Delaying * |
| *macloskeyi* | -0.220 | 0.001 | Advancing * |
| *nephrophylla* | -0.024 | 0.660 | Delaying * |
| *nuttallii* | 0.183 | 1.00E-04 | Delaying |
| *ocellata* | 0.394 | 0.005 | Delaying * |
| *odorata* | -0.022 | 0.852 | Advancing |
| *orbiculata* | -0.196 | 0.031 | Advancing |
| *palmata* | -0.381 | 2.11E-13 | Advancing * |
| *palustris* | -0.365 | 8.15E-14 | Advancing |
| ***pedata*** | **-0.452** | **8.60E-14** | Advancing * |
| *pedatifida* | -0.005 | 0.957 | Advancing |
| *pedunculata* | -0.083 | 0.122 | Delaying |
| *pinetorum* | 0.159 | 0.173 | Delaying |
| *praemorsa* | -0.137 | 0.001 | Delaying * |

| | | | |
|---|---|---|---|
| *primulifolia* | 0.012 | 0.858 | Advancing * |
| *pubescens* | -0.275 | 3.48E-05 | Advancing * |
| *purpurea* | -0.113 | 0.002 | Advancing * |
| *quercetorum* | 0.067 | 0.448 | Advancing |
| *rafinesquei* | -0.369 | 8.00E-11 | Advancing * |
| *rostrata* | -0.076 | 0.458 | Advancing * |
| *rotundifolia* | 0.051 | 0.657 | Delaying |
| *rugulosa* | -0.030 | 0.767 | Advancing |
| *sagittata* | -0.084 | 0.173 | Advancing * |
| *scopulorum* | 0.033 | 0.758 | Delaying |
| *sempervirens* | -0.023 | 0.781 | Delaying |
| *sheltonii* | -0.455 | 2.03E-06 | Delaying * |
| *sororia* | -0.293 | 4.77E-21 | Advancing * |
| *striata* | 0.140 | 0.031 | Advancing |
| *tricolor* | -0.157 | 0.227 | Advancing |
| *trinervata* | -0.182 | 0.129 | Delaying |
| *vallicola* | 0.249 | 9.95E-07 | Delaying |
| *walteri* | 0.263 | 0.030 | Advancing |

Table S3. ANCOVA results for the effect of species' traits on rate of flowering time shift for the subset of species that significantly shifted flowering since 1900 (as opposed to all species). Model: rate of flowering shift ~ range size + flower color + region (east/west/both) + origin (native/introduced) + mating system + life history (annual/perennial). * = significant

| | *Sum Sq* | *Df* | *F value* | *p-value* |
|---|---|---|---|---|
| *Range size* | 0.00205 | 1 | 3.5949 | 0.8168 |
| *Flower color* | 0.01666 | 2 | 0.0555 | 0.8002 |
| *Region* | 0.81213 | 2 | 11.0165 | 0.0010 * |
| *Origin* | 0.02891 | 1 | 0.7842 | 0.3890 |
| *Mating system* | 0.00523 | 1 | 0.1419 | 0.7114 |
| *Life history* | 0.00318 | 1 | 0.0862 | 0.7728 |

Table S4. Tukey posthoc contrast for species trait 'Region' for the subset of species that significantly shifted flowering since 1900 (as opposed to all species). The estimate shows the difference in flowering shift rate (days/year) between species in the three region categories (east/west/both).

| | *Estimate* | *Std. Error* | *t value* | *p-value* |
|---|---|---|---|---|
| *East - Both* | -0.14428 | 0.15572 | -0.927 | 0.623 |
| *West - Both* | 0.27585 | 0.15952 | 1.729 | 0.217 |
| *West - East* | 0.42013 | 0.08961 | 4.688 | <0.001 * |

Table S5. The model fits for both the simple (S) and environmental (E) models for all species and the results of the ANOVA test for model of best fit (Figure 3).

| Species | Adjusted $R^2$ (S) | P-value (S) | Adjusted $R^2$ (E) | P-value (E) | ANOVA P-value | Best fit model |
|---|---|---|---|---|---|---|
| *adunca* | 0.033 | 2.10E-15 | 0.290 | 6.16E-143 | 1.00E-130 | Environmental |
| *affinis* | 0.247 | 7.85E-14 | 0.380 | 3.87E-21 | 8.99E-10 | Environmental |
| *appalachiensis* | -0.014 | 5.70E-01 | -0.011 | 5.12E-01 | 3.43E-01 | None |
| *arvensis* | 0.487 | 3.42E-14 | 0.557 | 1.07E-15 | 5.99E-04 | Environmental |
| *bakeri* | 0.001 | 3.69E-01 | 0.087 | 6.96E-02 | 3.69E-02 | None |
| *beckwithii* | 0.073 | 7.75E-03 | 0.288 | 1.01E-07 | 7.03E-07 | Environmental |
| *blanda* | 0.209 | 1.07E-06 | 0.309 | 6.87E-09 | 2.65E-04 | Environmental |
| *canadensis* | 0.007 | 7.63E-03 | 0.145 | 8.11E-40 | 1.19E-39 | Environmental |
| *cucullata* | 0.311 | 1.72E-32 | 0.330 | 2.58E-33 | 1.56E-03 | Environmental |
| *cuneata* | 0.029 | 1.03E-01 | 0.365 | 1.05E-08 | 5.44E-09 | Environmental |
| *douglasii* | 0.110 | 7.64E-05 | 0.392 | 1.11E-15 | 4.19E-13 | Environmental |
| *eriocarpa* | 0.214 | 1.58E-18 | 0.272 | 9.64E-23 | 1.00E-06 | Environmental |
| *fimbriatula* | -0.024 | 8.74E-01 | 0.013 | 2.99E-01 | 1.02E-01 | None |
| *glabella* | 0.011 | 2.30E-02 | 0.396 | 1.89E-56 | 7.68E-57 | Environmental |
| *hallii* | 0.111 | 9.30E-03 | 0.561 | 3.34E-11 | 1.63E-10 | Environmental |
| *hastata* | 0.460 | 5.77E-19 | 0.477 | 1.66E-18 | 4.60E-02 | Environmental |
| *hirsutula* | 0.386 | 3.08E-08 | 0.437 | 1.78E-08 | 2.12E-02 | Environmental |
| *howellii* | -0.036 | 9.83E-01 | -0.072 | 9.92E-01 | 8.95E-01 | None |
| *labradorica* | 0.090 | 1.64E-07 | 0.720 | 3.22E-85 | 1.57E-80 | Environmental |
| *lanceolata* | -0.008 | 6.65E-01 | 0.003 | 3.59E-01 | 1.71E-01 | None |
| *lobata* | 0.050 | 1.83E-02 | 0.408 | 3.00E-13 | 5.96E-13 | Environmental |
| *macloskeyi* | 0.050 | 1.34E-03 | 0.344 | 5.75E-20 | 1.02E-18 | Environmental |
| *nephrophylla* | 0.029 | 3.31E-03 | 0.241 | 2.36E-19 | 1.66E-18 | Environmental |
| *nuttallii* | 0.019 | 5.68E-03 | 0.393 | 5.45E-48 | 1.04E-47 | Environmental |
| *ocellata* | 0.073 | 6.26E-02 | 0.116 | 4.84E-02 | 1.32E-01 | Simple |
| *odorata* | 0.067 | 3.28E-02 | 0.286 | 1.81E-05 | 4.13E-05 | Environmental |
| *orbiculata* | -0.005 | 4.87E-01 | 0.278 | 1.19E-08 | 1.29E-09 | Environmental |
| *palmata* | 0.518 | 1.72E-55 | 0.523 | 1.94E-54 | 5.96E-02 | Simple |
| *palustris* | -0.001 | 4.51E-01 | 0.281 | 1.72E-27 | 6.69E-29 | Environmental |
| *pedata* | 0.277 | 1.38E-18 | 0.299 | 7.13E-19 | 9.53E-03 | Environmental |
| *pedatifida* | 0.083 | 4.47E-03 | 0.335 | 3.67E-09 | 3.88E-08 | Environmental |
| *pedunculata* | 0.007 | 1.07E-01 | 0.090 | 4.11E-07 | 2.14E-07 | Environmental |
| *pinetorum* | -0.017 | 6.76E-01 | 0.557 | 1.33E-12 | 9.15E-14 | Environmental |
| *praemorsa* | 0.237 | 2.23E-38 | 0.523 | 4.32E-101 | 8.80E-66 | Environmental |
| *primulifolia* | 0.120 | 7.59E-08 | 0.117 | 7.90E-07 | 5.43E-01 | Simple |

| | | | | | | |
|---|---|---|---|---|---|---|
| *pubescens* | 0.405 | 1.67E-25 | 0.491 | 3.27E-31 | 2.11E-08 | Environmental |
| *purpurea* | 0.215 | 2.60E-40 | 0.457 | 9.84E-98 | 1.73E-60 | Environmental |
| *quercetorum* | 0.047 | 1.78E-02 | 0.308 | 3.01E-10 | 7.34E-10 | Environmental |
| *rafinesquei* | 0.425 | 1.00E-35 | 0.430 | 1.07E-34 | 9.61E-02 | Simple |
| *rostrata* | 0.540 | 3.70E-17 | 0.530 | 2.17E-15 | 9.95E-01 | Simple |
| *rotundifolia* | 0.131 | 1.77E-03 | 0.185 | 6.80E-04 | 3.45E-02 | Environmental |
| *rugulosa* | -0.016 | 8.24E-01 | 0.145 | 6.76E-04 | 8.43E-05 | Environmental |
| *sagittata* | 0.261 | 1.32E-18 | 0.263 | 1.80E-17 | 2.69E-01 | Simple |
| *scopulorum* | -0.019 | 8.40E-01 | 0.488 | 2.76E-12 | 1.47E-13 | Environmental |
| *sempervirens* | 0.008 | 2.11E-01 | 0.113 | 3.34E-04 | 1.46E-04 | Environmental |
| *sheltonii* | 0.044 | 4.22E-02 | 0.448 | 1.85E-12 | 1.76E-12 | Environmental |
| *sororia* | 0.305 | 5.90E-79 | 0.343 | 4.49E-89 | 3.09E-13 | Environmental |
| *striata* | 0.171 | 1.07E-10 | 0.197 | 3.08E-11 | 9.34E-03 | Environmental |
| *tricolor* | 0.254 | 7.51E-05 | 0.368 | 4.77E-06 | 3.67E-03 | Environmental |
| *trinervata* | -0.026 | 9.01E-01 | 0.115 | 1.49E-02 | 2.50E-03 | Environmental |
| *vallicola* | 0.016 | 1.84E-02 | 0.326 | 9.13E-32 | 7.67E-32 | Environmental |
| *walteri* | 0.021 | 1.87E-01 | 0.001 | 4.10E-01 | 7.21E-01 | None |

Table S6. Model results from ANCOVA evaluating whether rate of flowering shift (days/year) vary with different species traits. The results include the estimated marginal means (EMM), standard error (SE), degrees of freedom (df), and lower and upper confidence limits (CL) for each species' trait.

| | | *EMM* | *SE* | *Df* | *Lower CL* | *Upper CL* |
|---|---|---|---|---|---|---|
| *Range size* | | -0.188 | 0.0608 | 43 | -0.310 | -0.0651 |
| *Flower color* | *Violet* | -0.173 | 0.0691 | 43 | -0.313 | -0.0341 |
| | *White* | -0.165 | 0.0725 | 43 | -0.312 | -0.0193 |
| | *Yellow* | -0.224 | 0.0763 | 43 | -0.378 | -0.0705 |
| *Region* | *East* | -0.3016 | 0.0818 | 43 | -0.467 | -0.1366 |
| | *West* | -0.0379 | 0.0825 | 43 | -0.204 | 0.1285 |
| | *Both* | -0.2236 | 0.0755 | 43 | -0.376 | -0.0714 |
| *Origin* | *Native* | -0.148 | 0.0789 | 43 | -0.307 | 0.0113 |
| | *Introduced* | -0.228 | 0.1250 | 43 | -0.480 | 0.0248 |
| *Mating system* | *CHCL* | -0.171 | 0.0706 | 43 | -0.314 | -0.0288 |
| | *CH* | -0.204 | 0.0759 | 43 | -0.357 | -0.0512 |
| *Life history* | *Annual* | -0.254 | 0.1100 | 43 | -0.477 | -0.0314 |
| | *Perennial* | -0.121 | 0.0826 | 43 | -0.288 | 0.0451 |

Table S7. The change in co-flowering window (in days) for all pairwise species from historical (1900-1923) to current (2000-2023) time periods. On the diagonal, the values represent the change in flowering window (in days) for each species from the historical to current time periods, with positive values indicating a lengthening in flowering window and a negative values indicating a shortening of flowering window.

| | adunca | canadensis | cucullata | eriocarpa | glabella | macloskeyi | palustris | nephrophylla | nuttallii | palmata | pedata | praemorsa | pubescens | purpurea | sagittata | sororia |
|---|---|---|---|---|---|---|---|---|---|---|---|---|---|---|---|---|
| **adunca** | -39 | | | | | | | | | | | | | | | |
| **canadensis** | -27 | -16 | | | | | | | | | | | | | | |
| **cucullata** | -31 | -19 | 43 | | | | | | | | | | | | | |
| **eriocarpa** | -23 | -11 | 24 | 2 | | | | | | | | | | | | |
| **glabella** | -40 | -16 | 9 | -9 | -14 | | | | | | | | | | | |
| **macloskeyi** | -19 | -4 | -5 | -7 | 7 | 7 | | | | | | | | | | |
| **palustris** | -26 | -15 | -17 | -10 | -13 | -1 | -21 | | | | | | | | | |
| **nephrophylla** | -22 | -7 | -13 | -14 | -4 | -4 | -4 | -4 | | | | | | | | |
| **nuttallii** | -14 | 3 | -3 | -10 | 11 | 13 | 6 | 6 | 11 | | | | | | | |
| **palmata** | -20 | -19 | 41 | 20 | 3 | -11 | -18 | -19 | -12 | 42 | | | | | | |
| **pedata** | -32 | -20 | 27 | -1 | -21 | -23 | -19 | -23 | -25 | 24 | 9 | | | | | |
| **praemorsa** | -23 | -6 | -26 | -30 | -32 | -9 | -5 | -15 | -9 | -20 | -35 | -41 | | | | |
| **pubescens** | -18 | -6 | 26 | 22 | 22 | 8 | -4 | 0 | 7 | 18 | 8 | -13 | 35 | | | |
| **purpurea** | -35 | -17 | 5 | 0 | 1 | -6 | -14 | -6 | 11 | -1 | -16 | -20 | 18 | 1 | | |
| **sagittata** | -28 | -9 | 32 | 10 | 9 | 2 | -6 | -6 | -1 | 14 | -4 | -26 | 30 | 9 | 19 | |
| **sororia** | -29 | -5 | 59 | 7 | 9 | 6 | -2 | -2 | 3 | 42 | 11 | -26 | 37 | 13 | 19 | 60 |

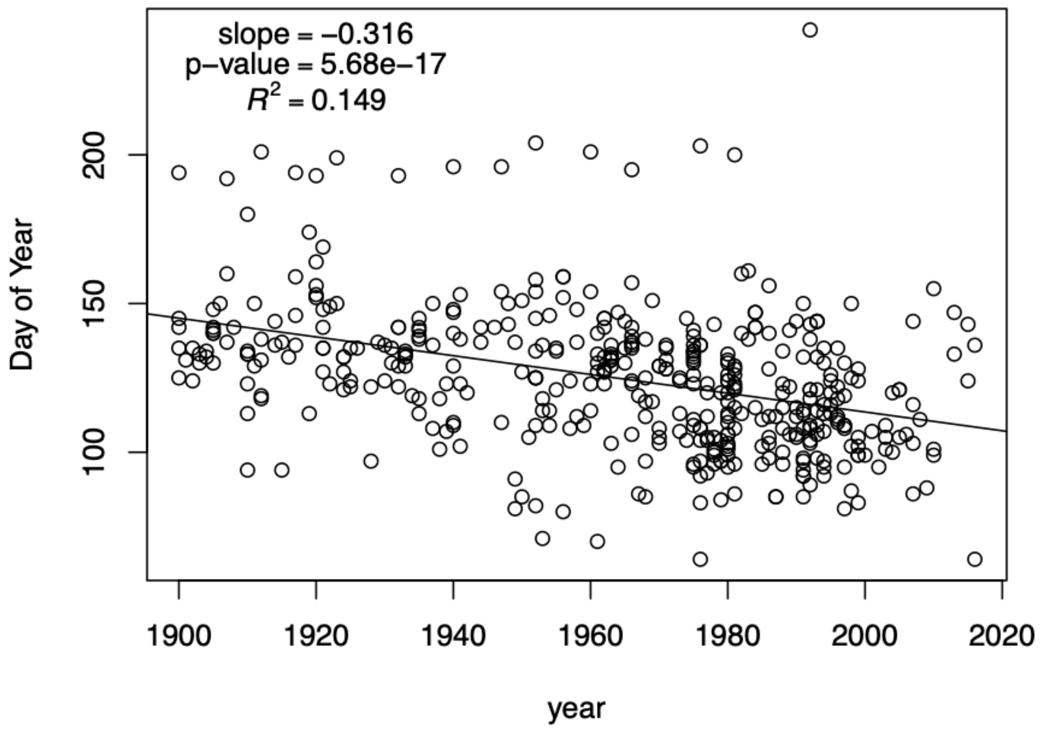


Figure S1. *Viola pubescens* flowering day over time. Dataset where we manually assessed the flowering status of all imaged herbarium specimens—a dataset that included both specimens annotated as flowering and specimens lacking this metadata (n = 431). The inferred flowering shift in this manually collected flowering dataset is -0.316 days/year, a shift that is statistically significant (p = 5.68e-17).

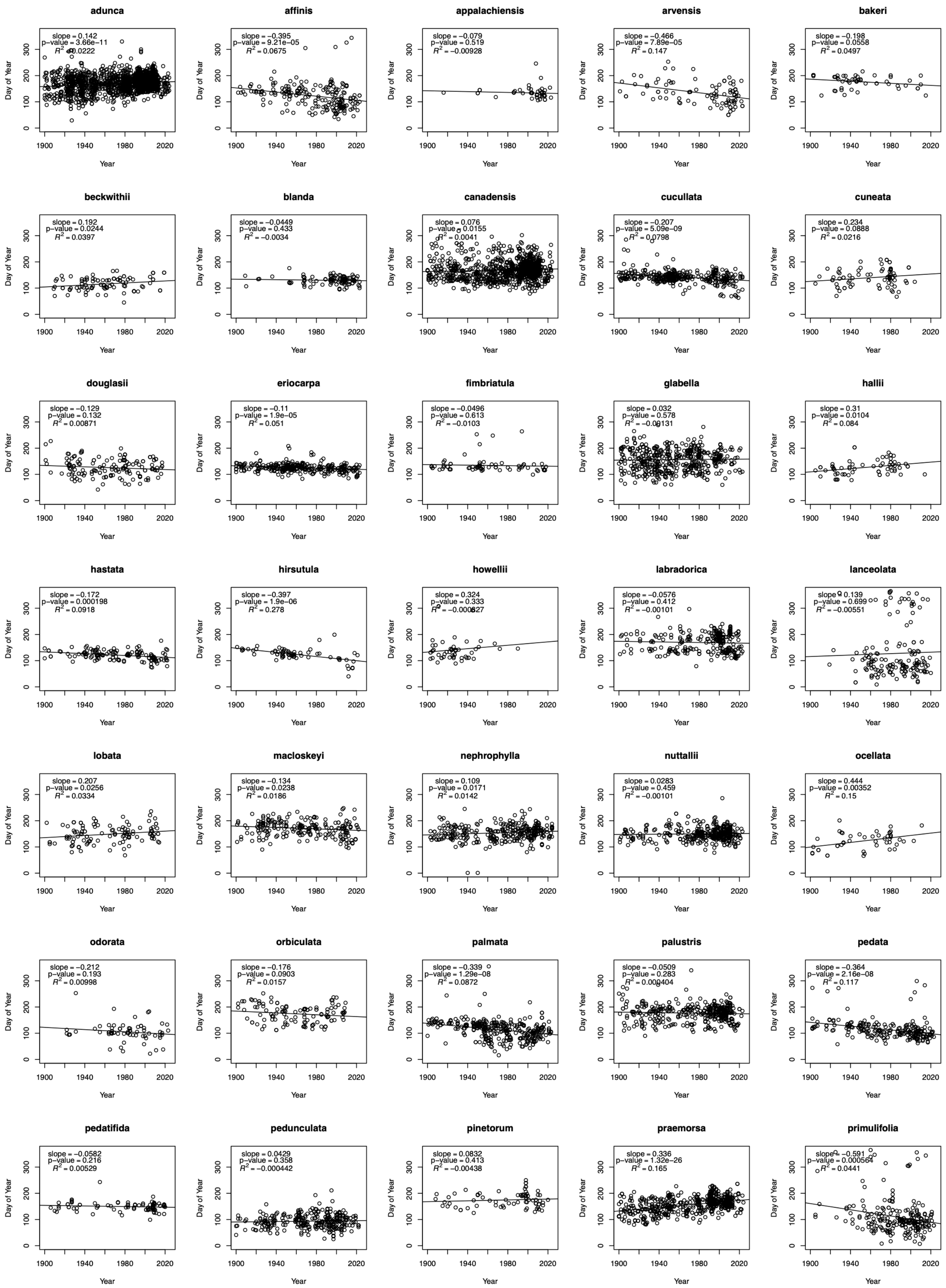
adunca
slope = 0.142
p−value = 3.66e−11
R^2 = 0.0222
affinis
slope = −0.395
p−value = 9.21e−05
R^2 = 0.0675
appalachiensis
slope = −0.079
p−value = 0.519
R^2 = −0.00928
arvensis
slope = −0.466
p−value = 7.89e−05
R^2 = 0.147
bakeri
slope = −0.198
p−value = 0.0558
R^2 = 0.0497
beckwithii
slope = 0.192
p−value = 0.0244
R^2 = 0.0397
blanda
slope = −0.0449
p−value = 0.433
R^2 = −0.0034
canadensis
slope = 0.076
p−value = 0.0155
R^2 = 0.0041
cucullata
slope = −0.207
p−value = 5.09e−09
R^2 = 0.0798
cuneata
slope = 0.234
p−value = 0.0888
R^2 = 0.0216
douglasii
slope = −0.129
p−value = 0.132
R^2 = 0.00871
eriocarpa
slope = −0.11
p−value = 1.9e−05
R^2 = 0.051
fimbriatula
slope = −0.0496
p−value = 0.613
R^2 = −0.0103
glabella
slope = 0.032
p−value = 0.578
R^2 = −0.00131
hallii
slope = 0.31
p−value = 0.0104
R^2 = 0.084
hastata
slope = −0.172
p−value = 0.000198
R^2 = 0.0918
hirsutula
slope = −0.397
p−value = 1.9e−06
R^2 = 0.278
howellii
slope = 0.324
p−value = 0.333
R^2 = −0.000827
labradorica
slope = −0.0576
p−value = 0.412
R^2 = −0.00101
lanceolata
slope = 0.139
p−value = 0.699
R^2 = −0.00551
lobata
slope = 0.207
p−value = 0.0256
R^2 = 0.0334
macloskeyi
slope = −0.134
p−value = 0.0238
R^2 = 0.0186
nephrophylla
slope = 0.109
p−value = 0.0171
R^2 = 0.0142
nuttallii
slope = 0.0283
p−value = 0.459
R^2 = −0.00101
ocellata
slope = 0.444
p−value = 0.00352
R^2 = 0.15
odorata
slope = −0.212
p−value = 0.193
R^2 = 0.00998
orbiculata
slope = −0.176
p−value = 0.0903
R^2 = 0.0157
palmata
slope = −0.339
p−value = 1.29e−08
R^2 = 0.0872
palustris
slope = −0.0509
p−value = 0.283
R^2 = 0.000404
pedata
slope = −0.364
p−value = 2.16e−08
R^2 = 0.117
pedatifida
slope = −0.0582
p−value = 0.216
R^2 = 0.00529
pedunculata
slope = 0.0429
p−value = 0.358
R^2 = −0.000442
pinetorum
slope = 0.0832
p−value = 0.413
R^2 = −0.00438
praemorsa
slope = 0.336
p−value = 1.32e−26
R^2 = 0.165
primulifolia
slope = −0.591
p−value = 0.000564
R^2 = 0.0441
Day of Year
Year

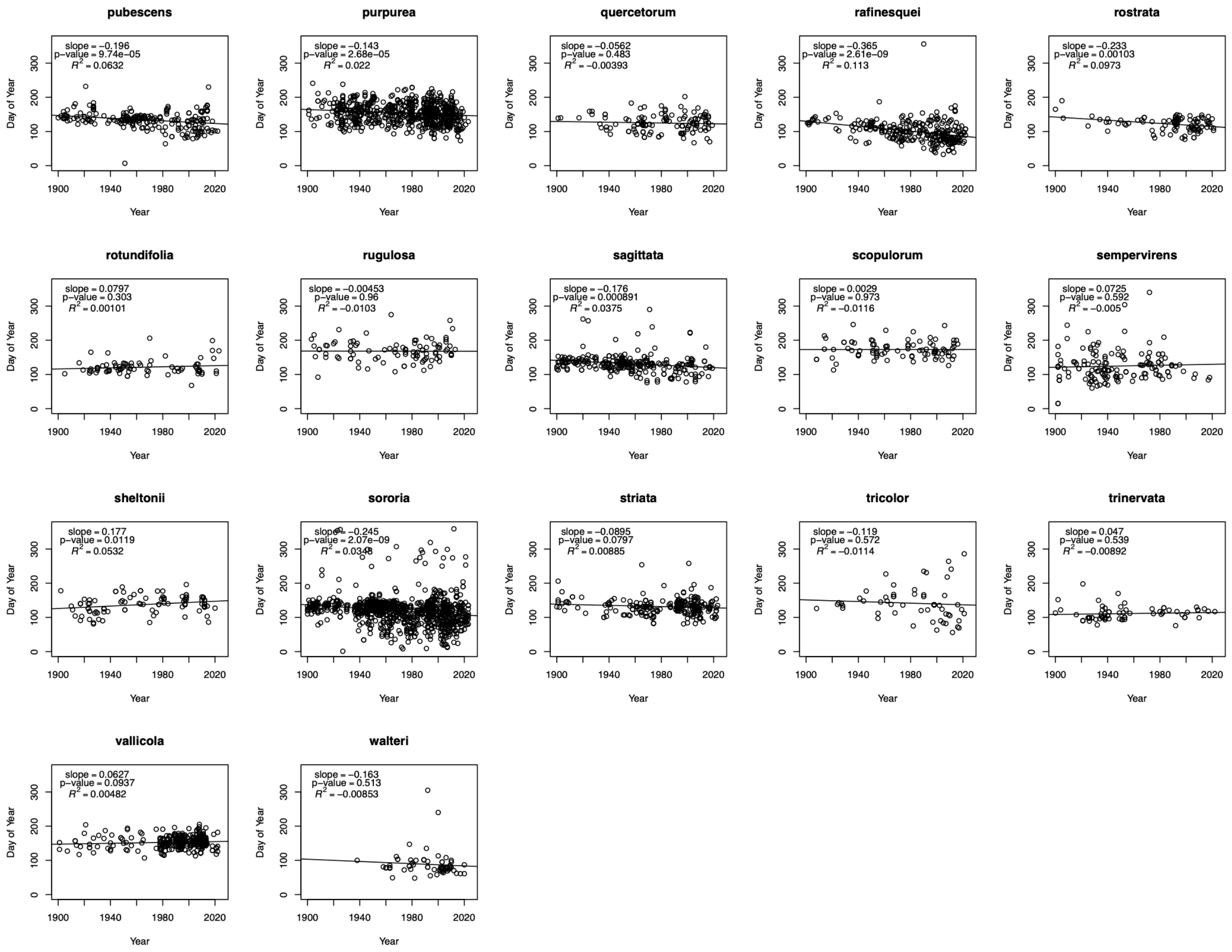


Figure S2. Linear model of flowering day as a function of year, displayed for all species individually. Each point within each plot represents one flowering occurrence record.

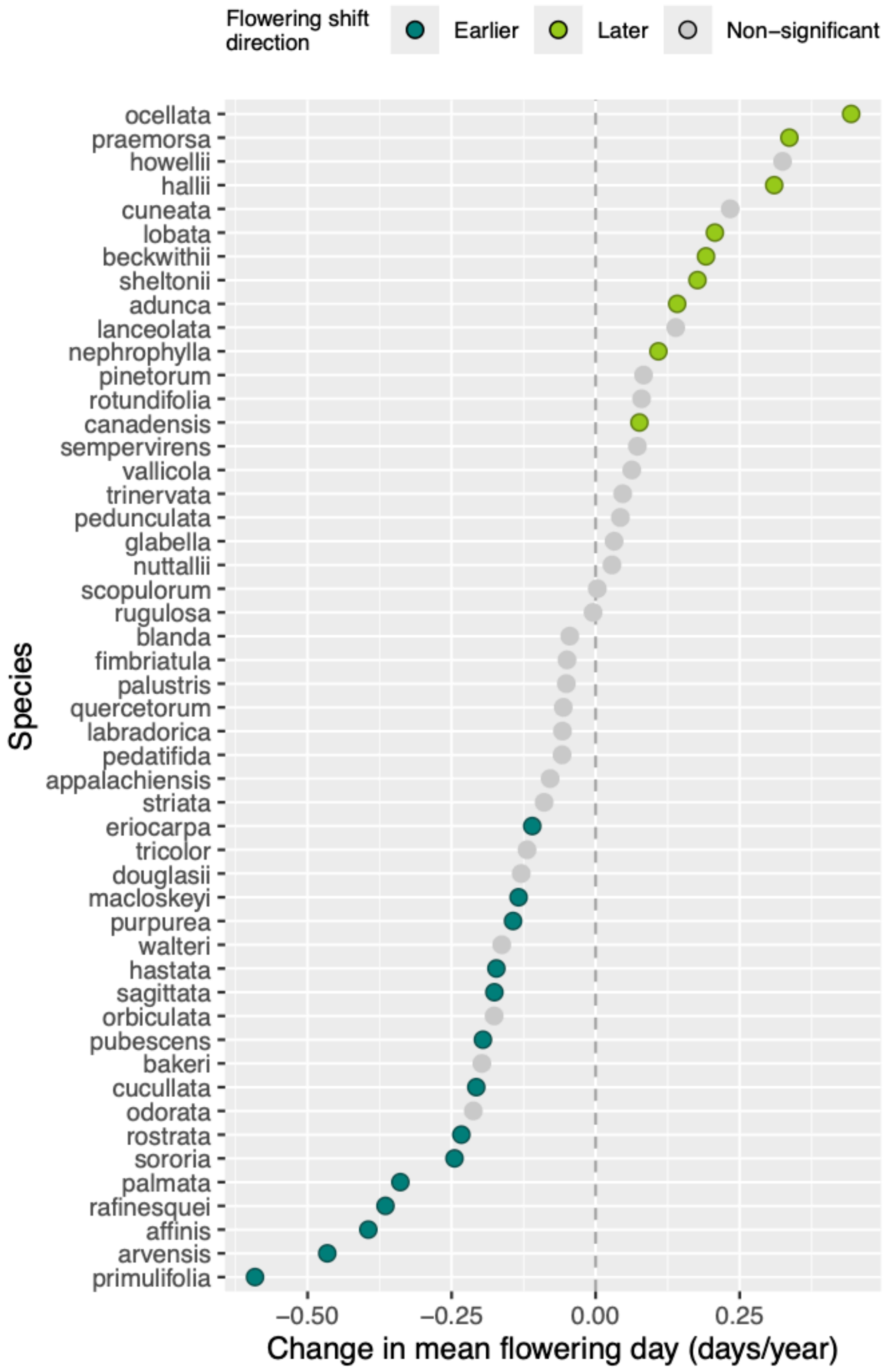


Figure S3. The rate of change in mean flowering day (days/year) for each species, based on samples from 1900-2024. Unlike the main text figure, this figure excludes species that had a year-latitude correlation coefficient r > 0.4 for advancing species and r < -0.4 for delaying species, a threshold that excludes species with high year-latitude correlation.

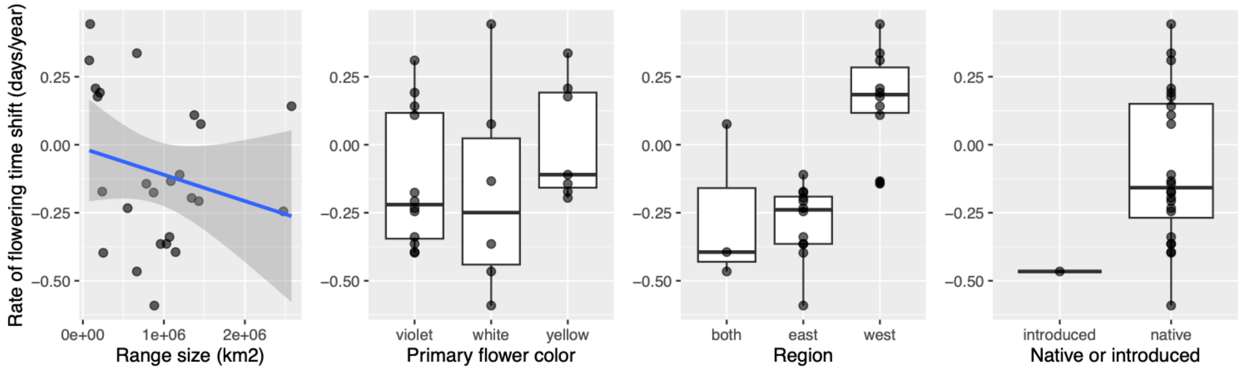


Figure S4. The rate of flowering phenology shifts (days/year) analyzed by A) range size (p = 0.817), B) flower color (p = 0.800) C) region (p = 0.001), and D) origin status (p = 0.389). Each point represents a species' rate of flowering change (from Figure 1). Unlike results reported in the main text (Figure 4), these analyses exclude species that did not significantly shift flowering phenology.

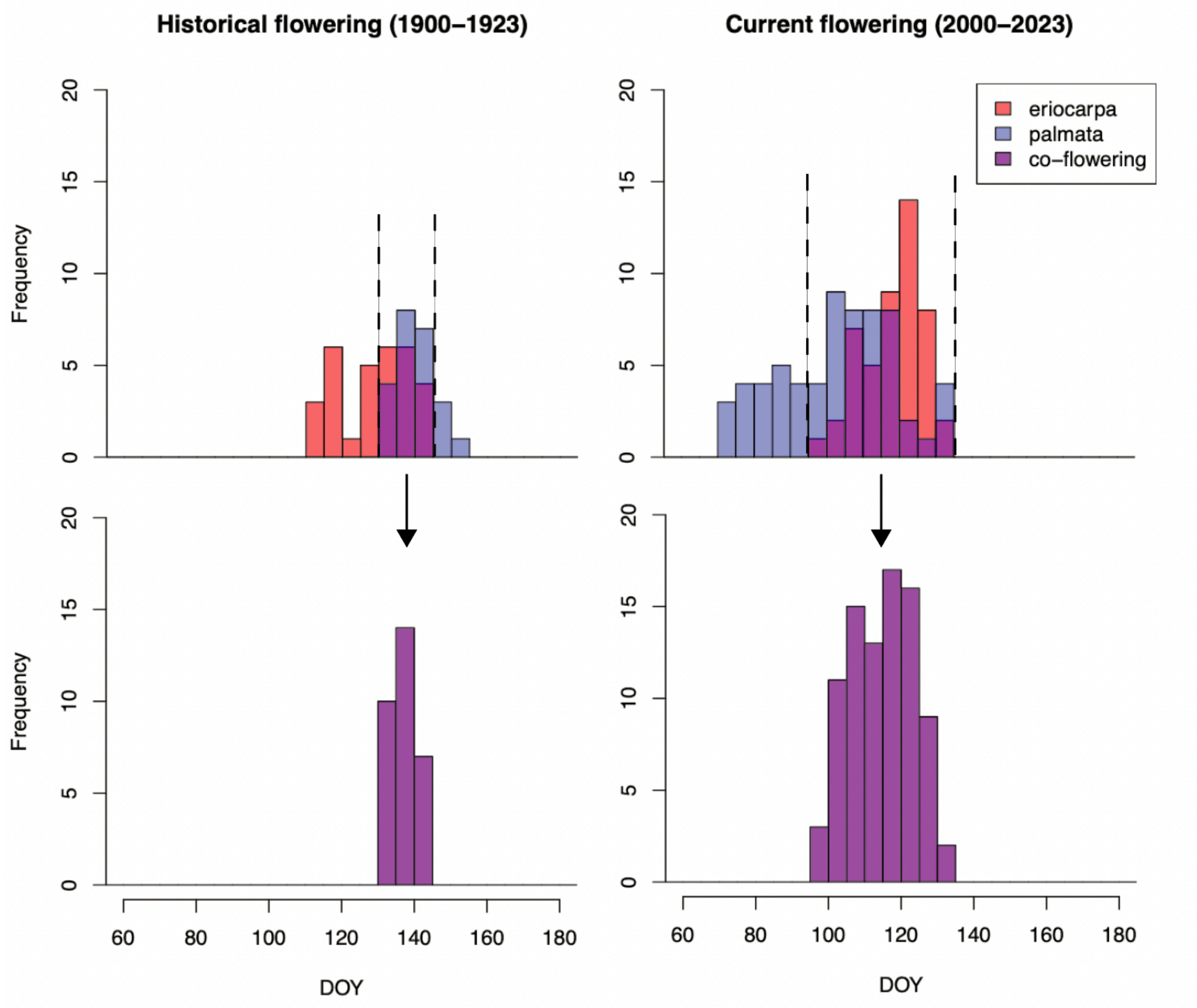

Figure S5. Example of how co-flowering distributions were defined. The top two panels show the distribution of flowering records over time (DOY) for two species (*Viola eriocarpa* and *Viola palmata*) and the overlap of these distr. The dashed lines define the start and end of the co-flowering window, when both species are flowering. The bottom two panels show the co-flowering distributions – all records from both species that fall within the co-flowering window.

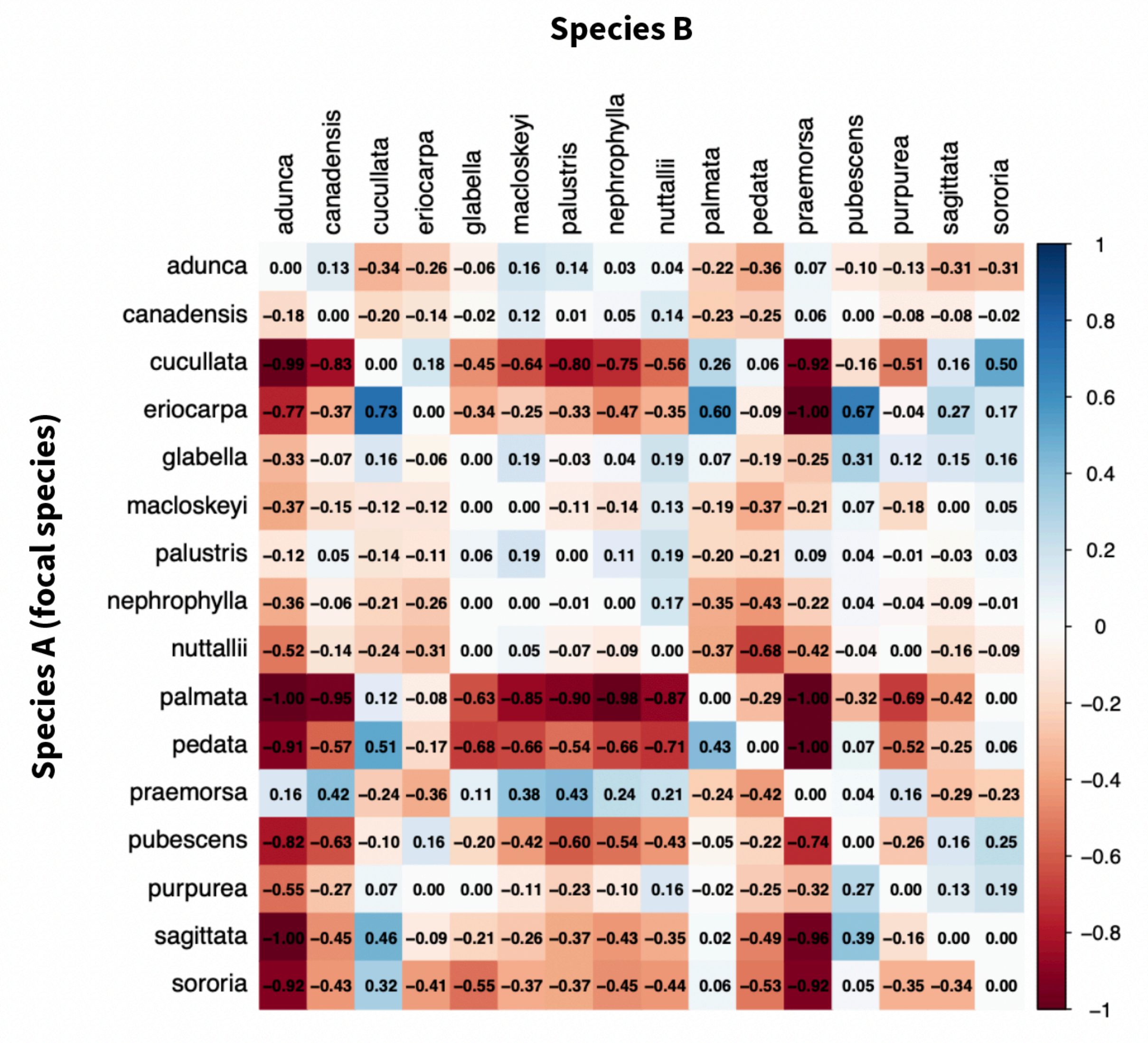


Figure S6. Co-flowering matrix that shows the change in flowering time overlap, measured by the difference in proportion of co-flowering out of total flowering window from 1900-1924 (past) to 2000-2024 (present) for all species pairs. The focal species is the denominator in this calculation (see Supplemental methods).

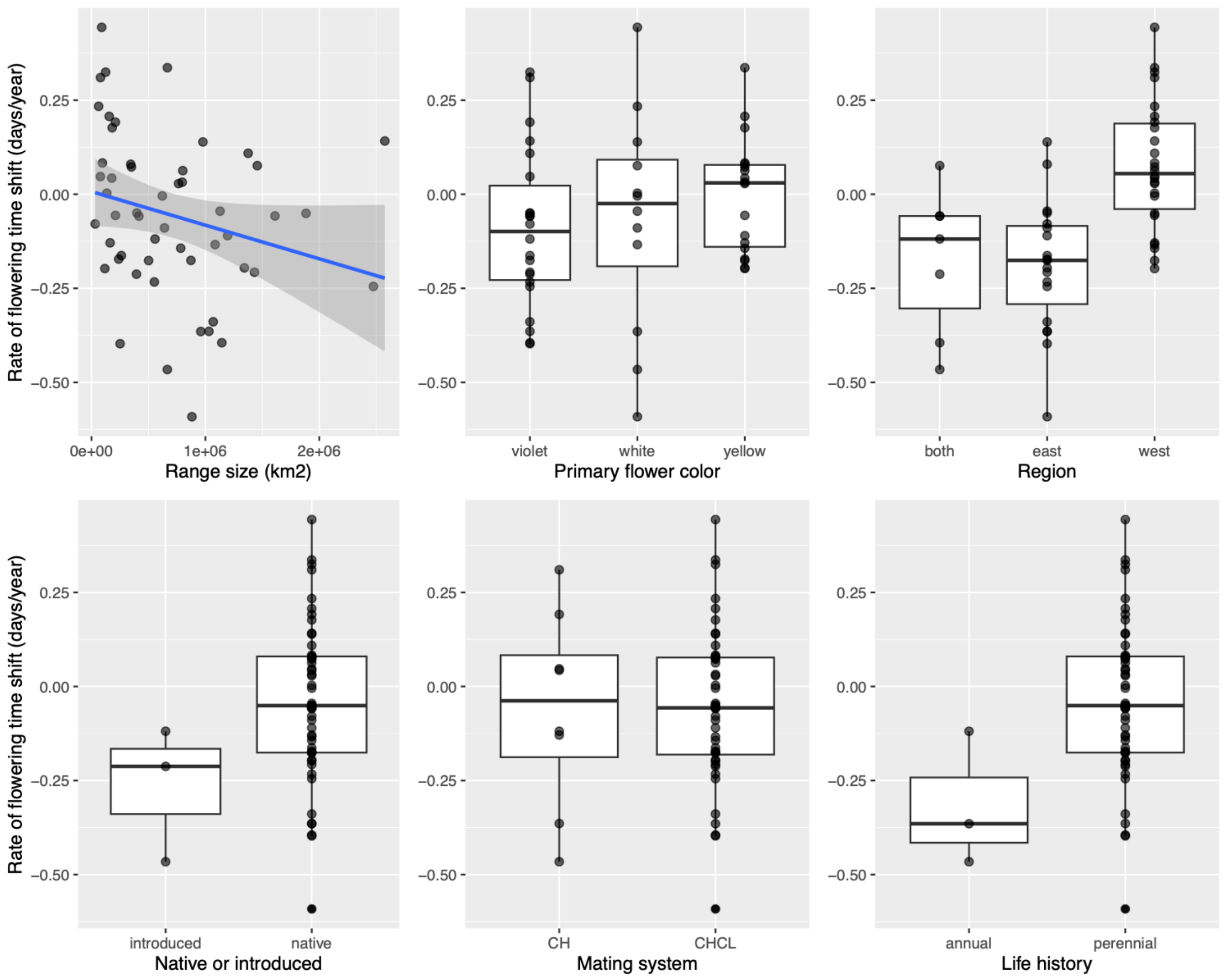


Figure S7. The rate of flowering phenology shifts (days/year) analyzed by A) range size ($p = 0.26$), B) flower color ($p = 0.65$) C) region ($p = 0.003$), D) origin status ($p = 0.64$), E) mating system ($p = 0.69$), and F) life history ($p = 0.39$). Each point represents a species' rate of flowering change (from Figure 1). Unlike results reported in the main text (Figure 4), these analyses include all species traits included in the model.

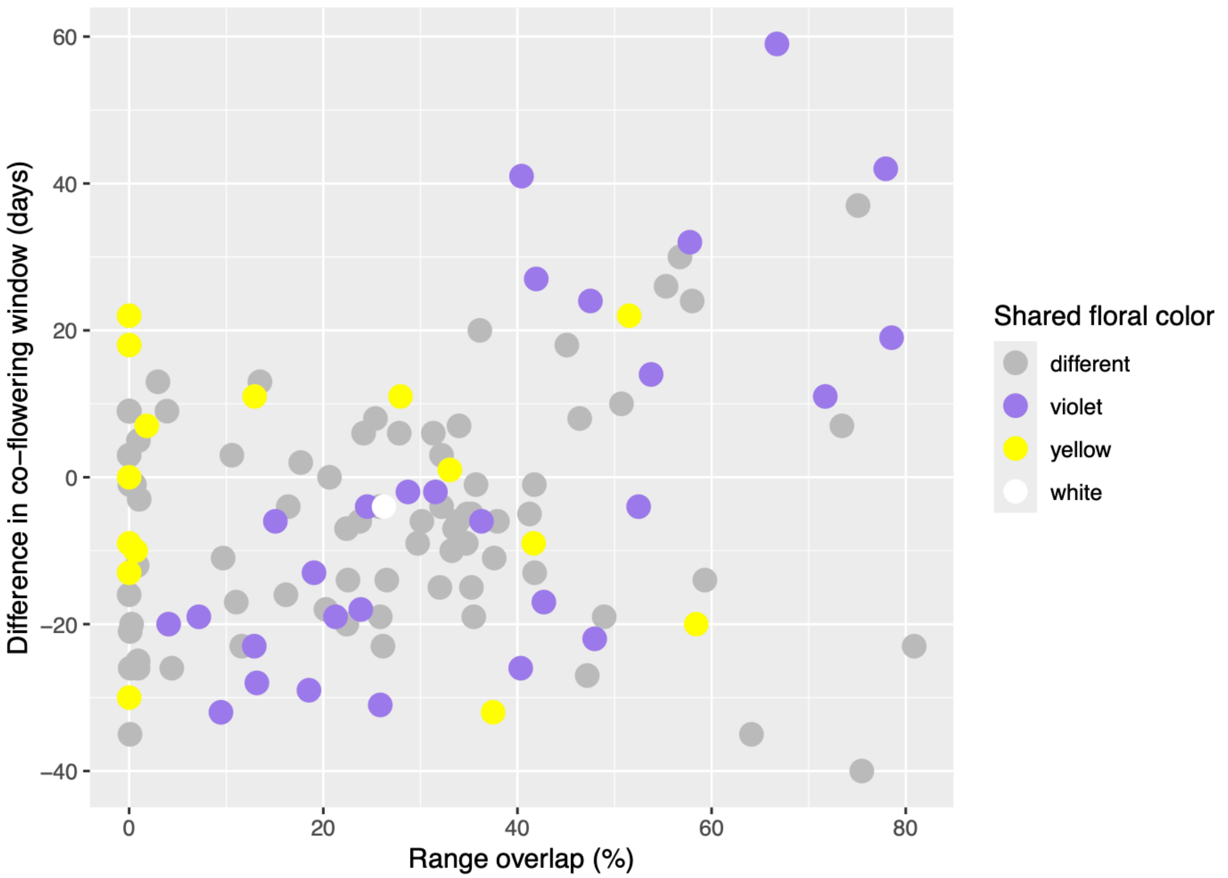


Figure S8. The relationship between range overlap percentage and the change in flowering overlap (days) from 1900-1924 (past) to 2000-2024 (present) for all species pairs (each point is a species pair). Points are colored by whether both species in a pair share a flower color (colored = shared flower color) or not (grey = different flower color).
